# Acoustic individual identification in a species of field cricket using deep learning

**DOI:** 10.1101/2025.05.24.655958

**Authors:** Emmanuel Kabuga, Diptarup Nandi, Stuart Burrell, Gciniwe Dlamini, Rohini Balakrishnan, Bubacarr Bah, Ian Durbach

## Abstract

Individual animal identification is essential for wildlife conservation and management, helping to estimate animal abundance and related parameters. We assessed the feasibility of identifying individual field crickets (*Plebeiogryllus guttiventris*) from their calls using deep learning. In a closed population of known individuals, the best models recognized crickets with up to 99.9% accuracy when trained and tested on calls from the same night, and 63.2% when tested on calls from nights not used in training. When matching pairs of calls, without knowledge of all individuals in the population, models identified pairs of calls from the same night with 97% accuracy, falling to 87.1% if calls were from different nights. Accuracy remained high when tested on individuals not observed during training (within night: 94.1%; across nights 78.8%). Pooling training data across multiple nights improved test accuracy for all models. Deep learning outperformed random forests, particularly on harder tasks, although both were able to discriminate individuals. Analysis of temperature effects on cricket calling behavior showed that higher temperatures were associated with shorter chirp durations and higher carrier frequencies. However, adjusting spectrograms for chirp duration alone, or for both duration and frequency, did not improve performance. The results are the first to demonstrate AIID using wild recordings of an insect species, and further highlight the potential of deep learning-based AIID for non-invasive monitoring of animal populations.

## 1 Introduction

Many ecological studies rely on being able to periodically detect, monitor or collect data from the same individual animal. Historically done using physical capture and marking, increasingly this is being done by manual or computer-assisted inspection of passive recordings made using camera or audio recorders, for species that are sufficiently visually or acoustically distinctive to be individually identifiable. Perhaps the most ubiquitous purpose for repeated identification of the same individual is to estimate the size of the population to which the individuals belong i.e. abundance, as well as related demographic parameters such as birth and mortality rates, using capture-recapture methods. Estimates of population size are a central component of evidence-based natural resource management (Johnston et al., 2015; Katzner, Ivy, Bragin, Milner-Gulland, & DeWoody, 2011), and are often imposed as legal requirements for the conservation and management of certain areas by national and regional governments (Evans, 2006).

However there are many other reasons for recording animal identity over time, for example studying the composition and dynamics of animal social networks (Schofield et al., 2019), or tracking changes in animal behaviour over time e.g. in response to environmental factors (Amin et al., 2022; Brunk, West, Peery, & Pidgeon, 2022). In short, individual animal identification is an essential preprocessing step for many ecological studies.

Resolving large quantities of passively recorded images or audio into individual animal identities is still largely done manually, but quickly becomes laborious, time consuming, and prone to error, particularly as data volumes increase (Crouse et al., 2017). The need for automated classifiers to facilitate and accelerate manual individual identification processes has long been recognised (Chen et al., 2020; Guo et al., 2020; Schneider, Taylor, & Kremer, 2020). Deep learning approaches have been used to successfully recognize individuals from photographic images, for an increasingly diverse set of species (e.g. dolphin (Blount et al., 2022); whale, fruit fly, and tiger (Schneider et al., 2020), giant panda (Chen et al., 2020); zebra (Dlamini & van Zyl, 2020), elephant (Korschens & Denzler, 2019), giraffe (Miele et al., 2021), lion (Dlamini & van Zyl, 2020), fish (Gómez-Vargas, Alonso-Fernández, Blanquero, & Antelo, 2023), seal, and toad (Kabuga et al., 2024)). Deep learning has also been widely used for species identification from acoustic recordings (e.g. Dufourq, Batist, Foquet, & Durbach, 2022; Eichinski, Alexander, Roe, Parsons, & Fuller, 2022; Gan et al., 2021; Yassir, Andaloussi, Ouchetto, Mamza, & Serghini, 2023).

Nevertheless, reports of individual identification from acoustic recordings have remained relatively rare (Linhart, Mahamoud-Issa, Stowell, & Blumstein, 2022) and focussed overwhelmingly on birds (Bedoya & Molles, 2021; Budka, Wojas, & Osiejuk, 2015; Stowell, Petrusková, Šálek, & Linhart, 2019) and large mammals with relatively distinctive vocalizations (African wild dog (Hartwig, 2005), grey wolf (Root-Gutteridge et al., 2014), African lion (Wijers et al., 2021), marmoset monkey (Oikarinen et al., 2019), bottlenose dolphin (Longden et al., 2020)). Acoustic individual identification (AIID) is only possible in presence of individual-specific acoustic signatures, which explains the focus on taxa with complex vocalizations, primarily birds and mammals (E. Knight et al., 2024; Stowell et al., 2019). In contrast, among the invertebrates, a large number of insects produce acoustic signals using the mechanism of stridulation. Many species of the insect order Orthoptera are particularly notable for their acoustic communication, with distinct species-specific acoustic signals (Gerhardt & Huber, 2002). AIID, however, has never been attempted in any of the Orthopteran species before. Few if any insects have ever been studied for individual identification using calls, even though some of these, such as field crickets, have served as model systems for studies on sexual selection and signal evolution for decades now. Successful AIID applications, particularly to these species with less-detailed call structures outside of mammal and avian taxa, would provide important indicators of the feasibility of automated approaches to monitor and study animal populations acoustically.

This study contributes to the emerging area of individual acoustic identification by developing models that discriminate between, and hence recognize, individuals of a species of field cricket *P. guttiventris* based on their acoustic calls (chirps). We compare the performance of traditional machine learning methods (random forests (RFs)) based on pre-selected features (Mel-frequency cepstral coefficients, MFCCs; temporal features such as call and inter-call durations) and deep learning algorithms (convolutional neural networks (CNNs)) trained on spectrogram images (Gupta, Kshirsagar, Zhong, Gholami, & Ferres, 2021; E. C. Knight, Poo Hernandez, Bayne, Bulitko, & Tucker, 2020) that provide a visual representation of how the frequency spectrum of a signal changes over time, thus capturing both temporal and spectral acoustic features. We assess the performance of these classifiers in both a “closed population” context (where the population size is assumed fixed and known for the duration of the survey and the task is to assign new calls to one of a set of known individual identities) and an “open population” context (where population size is unknown, and the task is to assess the similarity between two calls and decide if they belong to the same individual). Finally, we assess classifier performance when test calls are drawn from the same night(s) as those used to train the models, and in the more practically important setting when test calls are drawn from different nights. This is necessitated by the fact that field crickets, being insects, are poikilotherms, and the call features of individuals can change with temperature as well as diet across time (Ballesteros, Tan, & Robillard, 2022). Our results contribute to the limited literature on individual animal identification from acoustic recordings and provide a necessary first step that can potentially lead to a feasible approach for passive monitoring of insect communities.

## 2 Materials and Methods

### 2.1 Data description

The study analyzes an acoustic dataset of 137 audio recordings from 47 male crickets of the species *Plebeiogryllus guttiventris*. The recordings were made at a sampling frequency of 44.1 kHz, with an average duration of 86 seconds (ranging from 1 to 2 minutes). Each individual was recorded between two and five times over three nights in the village Ullodu (13^*°*^ 38^*′*^ N, 77^*°*^ 42^*′*^ E) in the state of Karnataka in India: all 47 individuals were recorded on the first night, 42 on the second, and 15 on the third, resulting in 54% of recordings on the first night, 35% on the second, and 11% on the third. The recordings, captured at a distance of 20 cm from the caller, are clean with minimal background noise. They include calling songs, used by males to attract females from afar, and courtship songs, used when females are nearby (Nandi & Balakrishnan, 2013). Each song consists of chirps with 1-6 syllables; there are very short periods of silence between syllables (≈ 5 ms), and longer intervals between chirps (≈ 45 ms). For further details on sampling design and recording, see Nandi and Balakrishnan (2013).

### 2.2 Cricket signal segmentation

#### 2.2.1 Chirp and fixed-length segment extraction

The first 10 seconds of each recording, consisting of the field researcher listing the recording identifier, were removed prior to analysis. Two acoustic-based segmentation algorithms were then implemented prior to AIID: a threshold-based chirp detector, and partitioning each recording into fixed-length segments.

Spectral gating (Sainburg, Thielk, & Gentner, 2020) was used for noise removal, followed by a low-pass filter to attenuate and smooth out any remaining high-frequency noise components. Amplitudes were standardized to lie between −1 and +1 to ensure consistency across all recordings.

We identified chirp start and end times as the earliest and latest times exceeding a signal-to-noise ratio amplitude of 0.1, with a maximum of 45 ms between threshold crossings for points to be in the same chirp. We chose the amplitude threshold based on ambient noise in the cricket signal, and we selected the between-chirp time threshold based on visual inspection of background noise intervals from several cricket recordings. We segmented the audio signal between start and end times to isolate detected chirps.

Although chirp and syllable-based acoustic segmentation is frequent in bio-acoustics, many classification applications use time intervals of fixed length (Dufourq et al., 2022; Lakshminarayanan, Raich, & Fern, 2009). As a result, we also extracted same-length segments by dividing each recording into time intervals (“segments”) of 1s duration, with 80% overlap between consecutive segments. Below, where the same processing step has been applied to both chirps and 1s segments, we refer to these collectively as “samples”.

#### 2.2.2 Manual postprocessing

The segmentation method exhibited near-perfect accuracy in extracting chirps from audio recordings. Upon visually inspecting the potential chirps generated by the segmentation algorithm, it accurately segmented chirps in 99.7% of cases (44,598 out of 44,730), with only 132 erroneous cases containing more than six syllables.

The generated chirp categories comprised 1,200 one-syllable, 4,880 two-syllable, 5,176 three-syllable, 8,638 four-syllable, 23,303 five-syllable, and 1,404 six-syllable chirps. Subsequent analysis focused exclusively on five-syllable chirps. Analyzing chirps with the same numbers of syllables is crucial for similarity learning methods that rely on visual comparisons, and additionally standardizes the number of temporal features extracted from chirps (see Section 2.3.1).

We manually assessed 51,776 1s segments produced by the segmentation method. Of these segments, 2,609 (5%) were background-noise segments with no entire chirps or syllables and thus were removed from the analysis. As a result, we only considered the remaining 49,167 segments belonging to 47 individuals for the subsequent analysis. Each individual produced 1,053 segments on average (min = 582 and max= 1,846). The variability in the number of segments primarily originates from more recordings being available for some individuals than others.

### 2.3 Cricket acoustic signal features

We employed features from five-syllable chirps and 1s segments to train RF and CNN models. While CNNs can automatically learn relevant features from images, random forests require preprocessed features, highlighting relevant patterns or characteristics of the data, as input. For CNN models, we derived spectrogram images from five-syllable chirps and 1s segments, as is standard practice (Stowell et al., 2019). RF models use temporal features and MFCCs for five-syllable chirps and MFCCs only for 1s segments (as these may contain multiple or partial chirps).

#### 2.3.1 Chirp temporal features and carrier frequency

Chirp temporal features include the duration of each syllable, the duration between each pair of syllables, the mean duration of syllables, the mean duration between syllables, and the chirp duration. We identified syllable start and end times using the approach described in Section 2.2.1. Start and end times are the earliest and latest times exceeding a signal-to-noise ratio amplitude of 0.1, with a maximum interval of 5.3 ms (235 samples) between threshold crossings for points to be considered part of the same syllable. The segmentation algorithm achieved very high accuracy in detecting syllables in five-syllable chirps. It correctly identified the number of syllables in 98.3% of the cases (22,918 out of 23,303). The remaining 385 chirps were discarded from the analysis. We computed the carrier frequency for the retained chirps. On average each individual produced 490 five-syllable chirps, with counts ranging from 20 to 1,350.

#### 2.3.2 Mel spectrograms

Spectrograms were generated for five-syllable chirps and 1s segments using a Hann window function with a length of 1,024 samples (23 ms), a hop length of 256 samples (5.8 ms, 75% overlap), and 128 mel-filter bands with centers evenly distributed between 4 and 8 kHz, as this range contained the entirety of *P. guttiventris* individual signals. We applied a 1024-point short-time Fourier transform to each windowed and filtered frame to extract a spectrogram depicting energy at various frequency bands over time (of 128 *×* 33 and 128 *×* 173 pixel resolution for five-syllable chirps and 1s segments, respectively). To ensure a uniform time resolution, five-syllable chirps were zero-padded to match the size of the longest five-syllable chirp prior to the construction of spectrograms. Figure 1 displays 1s sample spectrograms from two cricket individuals.

**Figure 1:**
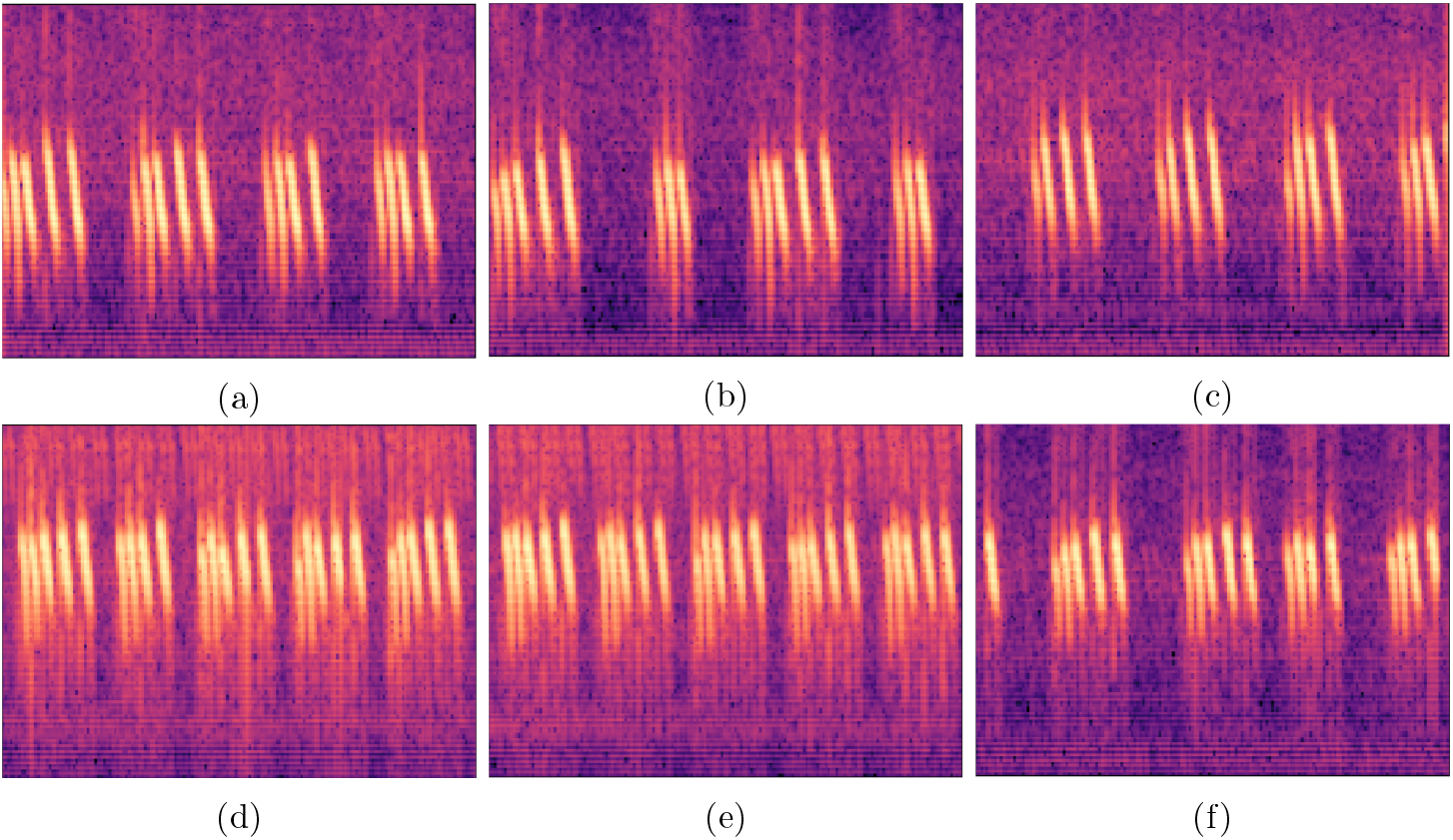
Samples of 1s spectrograms from two different crickets. Panels (a)–(c) correspond to the first cricket, and panels (d)–(f) to the second. Spectrograms (a) and (b) are taken from two different recordings of the first cricket recorded on the same night, while (c) was recorded on a different night. Similarly, (d) and (e) are from the same recording of the second cricket, recorded on a different night than (f).

#### 2.3.3 Mel-frequency cepstral coefficients

We extracted the first 20 MFCCs from each frame, using the same window length of 1,024. MFCC matrices derived from five-syllable chirps exhibited a variable number of frames due to the varying number of samples, whereas 1s segments produced MFCC matrices with consistent dimensions. Therefore, to create feature vectors of the same length, we summarized MFCC matrices by their means (20 entries) and standard deviations (20 entries) over time. Concatenating these two vectors produces a 40 *×* 1 input vector for both five-syllable chirps and 1s segments.

### 2.4 Standard classification and similarity learning approaches

#### 2.4.1 Standard classification neural networks

The “closed population” AIID task, where the number of calling individuals remains known and constant during the study period, is a standard multi-class classification task that can be addressed using standard CNNs (Figure 2, (1)). We used a model architecture consisting of six convolutional blocks (TABLE 1). Following standard CNN design practices (He, Zhang, Ren, & Sun, 2016), each of these blocks included 1–3 convolutional layers interspersed with max pooling and batch normalization layers. We used a larger filter size (7*×*7) in the first convolutional block to capture broad patterns and spatial relationships (e.g. edges, corners) within the input data (He et al., 2016; Schroff, Kalenichenko, & Philbin, 2015). The final four blocks each contains four residual neural network blocks (ResNet, He et al. (2016)) whose output includes both its input and the standard output, a reparameterization that has been found to improve performance in many applications (Schneider et al., 2020). We implemented a global max pooling layer after the last block to retain the maximum value from each feature map (Lin, Chen, & Yan, 2013).

**Table 1:** Foundational processing blocks of the CNN-based similarity learning model used to summarize a spectrogram into 512-vector embedding. The column “Output size” displays the output shape (height, width, channels) of the feature map at each processing block, while the column “Layer’s parameters” provides the number of filters, their size, and stride, respectively, applied at each layer designated by round brackets within a processing block. MP: Max-pooling, GMP: general max-pooling.

| Blocks | Output size | Layer’s parameters |
| --- | --- | --- |
| Block1 | (64, 87, 64) | [(64, 7, 2), (MP, 2, 2)] |
| Block2 | (32, 44, 64) | [(64, 3, 2), (64, 3, 2)] |
| Block3 | (16, 22, 128) | [(64, 1, 1), (64, 3, 1), (64, 1, 1)] × 4 |
| Block4 | (8, 11, 256) | [(64, 1, 1), (64, 3, 1), (64, 1, 1)] × 4 |
| Block5 | (4, 6, 384) | [(96, 1, 1), (96, 3, 1), (96, 1, 1)] × 4 |
| Block6 | (2, 3, 512) | [(128, 1, 1), (128, 3, 1), (128, 1, 1)] × 4 |
| GMP | (1, 1, 512) | — |

**Figure 2:**
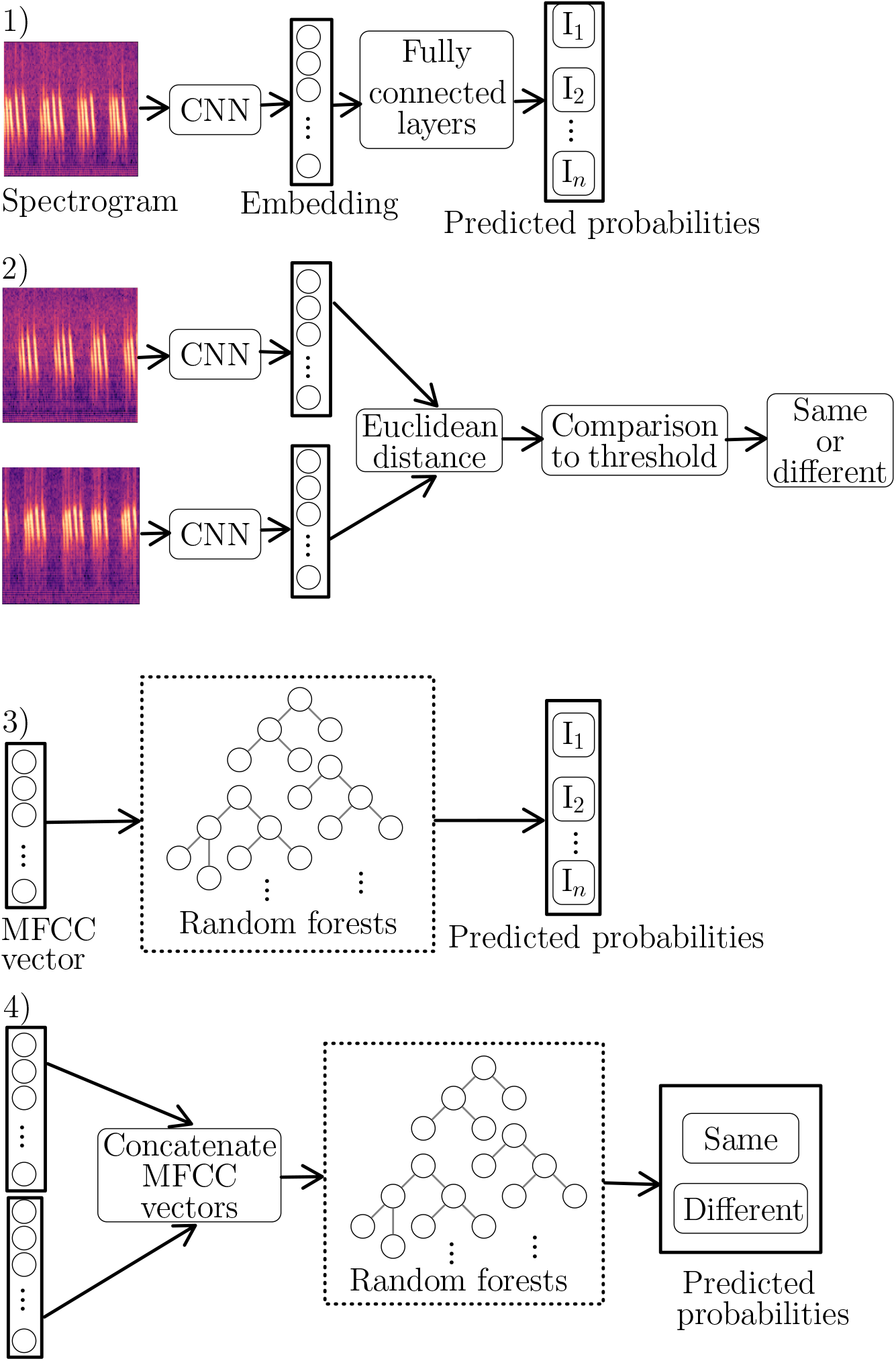
Summary of our methodologies. CNN and random forests standard classification models process input spectrograms (1) and MFCCs (2), respectively, to classify them into predefined cricket individuals. In contrast, similarity learning models (2 and 4) compare pairs of inputs to assess whether they belong to the same cricket individual based on their acoustic similarity.

Three fully connected layers using 256, 128, and 47 neurons then classify the learned embeddings to one of the 47 possible classes, each representing an individual. The first two fully connected layers use the ReLU non-linear action function. The last layer applies a soft-max activation function to convert predictions into probabilities.

Selection of hyperparameters such as the number and composition of convolutional blocks, number of filters and their size, activation functions, and learning rate is ideally by optimization over a large set of candidate parameter values. While this is important to optimize performance, it is very computationally expensive, and where good (rather than the best) performance is acceptable, many combinations likely offer comparable results. Because our intention is to demonstrate that *P. guttiventris* individuals can be accurately individually identified, rather than to maximize that accuracy, we adopted standard values for hyperparameters, such as 64 to 128 filters with sizes of 3 *×* 3 and 7 *×* 7, the pooling window size of 2 *×* 2, the use of the ReLU (rectified linear unit) activation function, and a stride of 1 or 2 (TABLE 1).

#### 2.4.2 Similarity learning neural networks

For “open population” AIID, the number of calling individuals is assumed unknown and the core task becomes to recognize whether two calls are from the same individual or not. We used triplet neural networks (Hermans, Beyer, & Leibe, 2017; Schroff et al., 2015) to perform this task. The triplet neural network is a two-phase model. During the embedding learning phase, it uses three CNNs with shared parameter values to summarize triplets of input images, with the aim of minimizing distances between pairs from the same individual and maximizing distances between pairs from different individuals within the embedding space. In the identification phase, two of the three CNNs summarize a pair of input images into two embeddings, compute the distance between them, and decide whether they belong to the same individual based on a distance threshold (Figure 2, (2)). We used the same CNN architecture and hyperparameter settings described in the previous section (TABLE 1).

#### 2.4.3 Random forests

To benchmark the performance of the CNN models, we trained random forest models to either classify each input vector into one of the 47 individual categories (closed population, Figure 2, (3)) or to produce a binary classification of whether the concatenated input vector from two individuals was from the same individual or not (open population, Figure 2, (4)). Hyperparameters – the number of trees used and the number of features available at each split – were selected by 3-fold cross validation over a grid of candidate values (number of trees: {200, 400, 600, 1000}; number of features: {5, 6, 7, 10}).

### 2.5 Construction of training, validation, and testing data

To assess the extent to which classifier performance degraded if tested on data from (a) different nights than those used to train the model; (b) different individuals than those used to train the model (open populations only), we constructed a number of partitions of the data into training, validation, and testing samples.

We trained models on recordings from either (a) one night (always night 1) or (b) two nights (always night 1 & 2), and tested the trained models on recordings made from (i) the same nights as those used in training, or (ii) on recordings made on different nights. For both closed- and open-population classifiers we begin by allocating samples to training, validation, and test sets. Open-population classifiers require the additional step of forming pairs or triplets of samples, and since creating all possible pairs or triplets is infeasible some sampling strategy is required. We describe this in Sections 2.5.3 and 2.5.4. In the following it will be useful to refer to *n*_*i*_ as an arbitrary sample from night *i*, (*n*_*i*_, *n*_*j*_) be a pair of samples where one is from night *i* and one is from night *j*, and 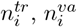, and 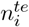 as a sample from night *i* allocated to training, validation, or test set respectively (with similar superscripts used for pairs).

#### 2.5.1 Allocation of samples

Where the same individuals were used for training and testing, we randomly designated 60% of each individual’s samples in each of night 1 and 2 as training samples, with the remaining 40% of samples in each night divided equally between validation and test sets. No training is done using samples from night 3 and so all of these are designated as test samples (TABLE 2 (a)).

**Table 2:** Data partition for the case where test samples are from the same individuals as training data or new individuals.

| Data | Train | Validation | Test | Total |
| --- | --- | --- | --- | --- |
| (a) Same individuals |  |  |  |  |
| 1s segments |  |  |  |  |
| Night 1 ($n_1$) | $n_1^{tr} = 15,608$ | $n_1^{va} = 5,208$ | $n_1^{te} = 5,238$ | 26,054 |
| Night 2 ($n_2$) | $n_2^{tr} = 10,663$ | $n_2^{va} = 3,560$ | $n_2^{te} = 3,580$ | 17,803 |
| Night 3 ($n_3$) | — | — | $n_3^{te} = 5,310$ | 5,310 |
| Total | 26,271 | 8,768 | 14,128 | 49,167 |
| Five-syllable segments |  |  |  |  |
| Night 1 ($n_1$) | $n_1^{tr} = 7,077$ | $n_1^{va} = 2,366$ | $n_1^{te} = 2,395$ | 11,838 |
| Night 2 ($n_2$) | $n_2^{tr} = 4,865$ | $n_2^{va} = 1,629$ | $n_2^{te} = 1,650$ | 8,144 |
| Night 3 ($n_3$) | — | — | $n_3^{te} = 2,936$ | 2,936 |
| Total | 11,942 | 3,995 | 6981 | 22,918 |
| (b) New individuals |  |  |  |  |
| 1s segments |  |  |  |  |
| Night 1 ($n_1$) | $n_1^{tr} = 14,583$ | $n_1^{va} = 6,293$ | $n_1^{te} = 5,178$ | 26,054 |
| Night 2 ($n_2$) | $n_2^{tr} = 8,897$ | $n_2^{va} = 3,814$ | $n_2^{te} = 5,092$ | 17,803 |
| Night 3 ($n_3$) | — | — | $n_3^{te} = 5,310$ | 5,310 |
| Total | 23,480 | 10,107 | 15,580 | 49,167 |
| Five-syllable segments |  |  |  |  |
| Night 1 ($n_1$) | $n_1^{tr} = 6,622$ | $n_1^{va} = 2,877$ | $n_1^{te} = 2,339$ | 11,838 |
| Night 2 ($n_2$) | $n_2^{tr} = 3,991$ | $n_2^{va} = 1,733$ | $n_2^{te} = 2,420$ | 8,144 |
| Night 3 ($n_3$) | — | — | $n_3^{te} = 2,936$ | 2,936 |
| Total | 10,613 | 4,610 | 7,695 | 22,918 |

Where the test set included new individuals not previously seen during training, we set aside audio recordings from 10 individuals across night 1, 2, and 3 for testing. Additionally, all samples from night 3 were included in the test set, resulting in 15 test individuals from night 3. We randomly allocated 70% of the remaining samples in each night to the training set and 30% to the validation set (TABLE 2 (b)).

#### 2.5.2 Closed population datasets

For the closed-population task, we created two training datasets (each with a corresponding validation dataset) and three test datasets for both 1s segments and five-syllable chirps (TABLE 2 (a)). For example, in the case of 1s segments, training datasets used either (1) training samples from night 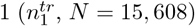 or (2) training samples from night 1 and 2(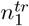 and 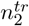, *N* = 26,271). Test datasets used test samples from (1) night 1 only 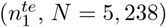, (2) night 2 only 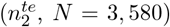, or (3) night 3 only 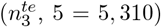. We evaluated the performance of models trained on each training set on each of the three test sets. The same procedure was applied to five-syllable segments.

#### 2.5.3 Pair construction

For the open-population task, we created two training datasets (each with a corresponding validation dataset) and six test datasets. Training datasets used either (1) pairs of training samples from night 1 only 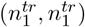 or (2) pairs of training samples drawn randomly after pooling training samples from night 1 and night 2—these pairs may be 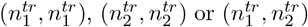. Test datasets used either pairs of samples (1) both from night 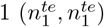, (2) both from night 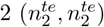, (3) both from night 3 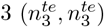, or (4–6) by pairing test samples in different nights 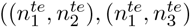, and 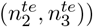. For analyses using the same individuals in model training and testing, pairs are formed from the data partitions in TABLE 2 (a). For analyses in which test datasets include new individuals not seen during training, pairs are chosen from TABLE 2 (b). Again, we evaluated the performance of models trained on either training set on each of the six test sets.

Matching training pairs were created by pairing each sample with another sample from the same individual drawn randomly without replacement, ensuring that each sample appeared exactly twice. Non-matching pairs were created similarly by pairing each sample with another sample from a different individual, again sampling without replacement. Aggregating matching and non-matching pairs over all individuals produced a balanced training dataset, twice the size of the original sample set used to produce it, in which each individual contributed an equal number of samples—two in matching pairs and two in non-matching pairs (e.g. *N* = 31, 216 and *N* = 52, 542 for training datasets (1) and (2) aforementioned, for the case where training and test sets are from the same individuals). Validation pairs were created similarly. Test pairs (matching or non-matching) were created by randomly sampling 5,000 pairs from the set of all possible (matching or non-matching) pairs.

#### 2.5.4 Triplet construction

During the embedding learning phase, triplet construction followed a strategy similar to that used in pair generation. Training datasets were composed either of (1) triplets exclusively from night 1—denoted as 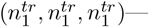 or (2) triplets from both night 1 and night 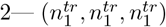 and 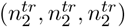, and six mixed-case scenarios allowing combinations of samples from both nights. In every triplet (*R, P, N*), the first element was always the reference example *R*, followed by the positive *P* and negative *N* examples. The same procedure was applied to validation triplets construction.

Triplets were created using the same online construction of semi-hard triplets used in Kabuga et al. (2024). The core idea in using a triplet input (Schroff et al., 2015) is that the model adjusts its parameters during training so that the similarity between the matching pair of samples in the triplet (as measured by inverse distance) is greater than the similarity between the non-matching pair of samples. For efficient model training, triplets in which this relationship does not already hold should be preferentially sampled, as compliant triplets make little or no contribution to the loss function. In particular, given a reference example this means selecting a dissimilar positive example and a similar negative example to make up the triplet. Schroff et al. (2015) showed that performance is best if, rather than selecting positive and negative examples that are respectively maximally dissimilar and similar to the reference example, triplets are constructed so that the reference example is slightly (i.e. within a specified margin) more similar to the positive example than the negative example. These are called semi-hard triplets, ecological applications of which are e.g. Kabuga et al. (2024); Moskvyak, Maire, Dayoub, Armstrong, and Baktashmotlagh (2021); Schneider et al. (2020).

Assessing the similarity between samples requires knowledge of the model’s current state, motivating an online approach to training in which triplets are created within each batch rather than *a priori*, as is done in “offline” mode (Schroff et al., 2015). We used batches of 128 spectrograms randomly drawn from at least 8 individuals. We then used the current state of the triplet neural network to summarize the spectrograms into their corresponding embeddings. Subsequently, we generated all possible triplets but used only semi-hard triplets for parameter updates during training. No test triplets were constructed; instead, the trained model was evaluated on the identification task using each of the six test pairs generated in Section 2.5.3, following the evaluation procedure outlined below in Section 2.7.

### 2.6 Model training

We trained CNN models for 30 epochs to minimize the semi-hard triplet loss for similarity learning models and categorical cross-entropy loss for standard classification models. We iteratively updated parameters using the Adam optimization algorithm with a learning rate ranging from 10^−3^ to 10^−5^. Initially set at 10^−3^, the learning rate was reduced by a factor of 0.5 whenever the validation performance did not improve over 5 epochs. Upon reaching the lower limit of 10^−5^, the model executed the remaining epochs using the minimum learning rate.

### 2.7 Optimized distance threshold and model evaluation

In the identification phase, following the embedding learning stage, an optimized distance threshold was used to separate matching from non-matching test pairs. This threshold was selected to maximize the F1-score (the harmonic mean of precision and recall) on validation pairs. A test pair was classified as a match if the Euclidean distance between its embeddings was below the threshold; otherwise, it was considered a non-match. Model performance was evaluated using four standard classification metrics for similarity learning models: accuracy (the proportion of correctly identified matches and non-matches), precision (the proportion of true matches among all predicted matches), recall (the proportion of actual matches that were correctly identified), and F1-score. For standard classification models, only overall accuracy was used.

### 2.8 Temperature analysis

Temperature significantly influences the acoustic properties of calls for many cricket species (e.g. *Gryllus integer* Martin, Gray, and Cade (2000); *Gryllus fultoni* and *Gryllus vernalis* Jang and Gerhardt (2007)), and a dependence between temperature and chirp duration, chirp period, syllable duration, and carrier frequency has already been shown for the set of *P. guttiventris* recordings we use Nandi and Balakrishnan (2013). Temperature-induced changes to spectrograms pose challenges for AIID and can be expected to reduce model performance unless accounted for. While specific methodologies vary, a general strategy is to normalize the temporal parameters of calls to a standard reference temperature before spectrogram construction, allowing for consistent comparisons across recordings to be made.

To assess the effect of temperature on our results, we tested for a relationship between temperature difference and the probability that a pair of 1s call segments from the same individual was more likely to be misclassified i.e. as two different individuals by our best CNN models. As there was no within-night variation in temperature for any individual, we used only pairs of calls recorded on different nights. We used binomial generalized linear mixed models (GLMMs) with individual as a random effect for individual to control for individual-specific differences in classification accuracy, implemented in the R package *lme4*.

To explore whether temperature normalization of spectrograms improved model performance, we estimated the relationship between two key temporal features, chirp duration and carrier frequency, and temperature, using GLMMs with a gaussian error structure and individual as a random effect. We then used these relationships to adjust five-syllable chirp duration and carrier frequency to a standard temperature of 29.1^*°*^*C* (the mean temperature across recordings) i.e. *f* (29.1) = *f* (*τ*) + *β*_*f*_ (29.1 − *τ*), where *f* (*t*) is the value of the temporal feature at temperature *t, τ* is the temperature the call was recorded at, and *β*_*f*_ is the temperature coefficient in the regression of the temporal feature on temperature. Time- and pitch-stretching corrections were implemented using the `time_stretch` and `resample` functions in the Python library *librosa*. We then constructed sets of spectrograms from chirps (a) with no adjustment for temperature, (b) adjusting for temperature effects on chirp duration, (c) adjusting for temperature effects on both chirp duration and carrier frequency. Below we first present results from unadjusted spectrograms, reporting the results on adjusted spectrograms for temperature effects in Section 3.4.

## 3 Results

### 3.1 Closed-population individual classification accuracy

The best-performing models recognized cricket individuals from the same nights they were trained on with high accuracy and crickets from other nights with moderate accuracy. Because results were very similar on 1s segments and five-syllable chirps, we only present the ones on 1s segments in the main body of the manuscript and the rest in Supplementary Material A.1.

The best CNN and RF models trained on segments from the first night achieved high accuracy in assigning segments from the same night, *n*_1_, to the cricket emetting the call, with average accuracies of 99.9% and 98.9%, respectively (Figure 3). Even when tested on segments from different nights as training data, *n*_2_ and *n*_3_, the models achieved substantially better results than random guessing classifiers, with average accuracies of 30.8% for CNN and 18.6% for RF. For example, assuming all individuals have the same number of test samples, a naive benchmark classifier allotting all samples to the majority class would achieve an accuracy of 1/47 = 2.1%. Likewise, the best-performing CNN and RF models trained on pooled segments from the first and second nights achieved high accuracy in recognizing cricket individuals from these nights, *n*_1_ and *n*_2_, with average accuracies of 99.8% and 98.7%, respectively (Figure 3). Their recognition accuracy over segments from a different night as training data, *n*_3_, increased to 63.2% and 30.0%, respectively. Overall, incorporating two nights into the training dataset significantly increased the classification accuracy on segments from different nights by 32.4% for the CNN models (from 30.8% to 63.2%) and 11.4% for the RF models (from 18.6% to 30.0%), while maintaining the classification accuracy on segments from the same nights as training data very high for both CNN and RF.

**Figure 3:**
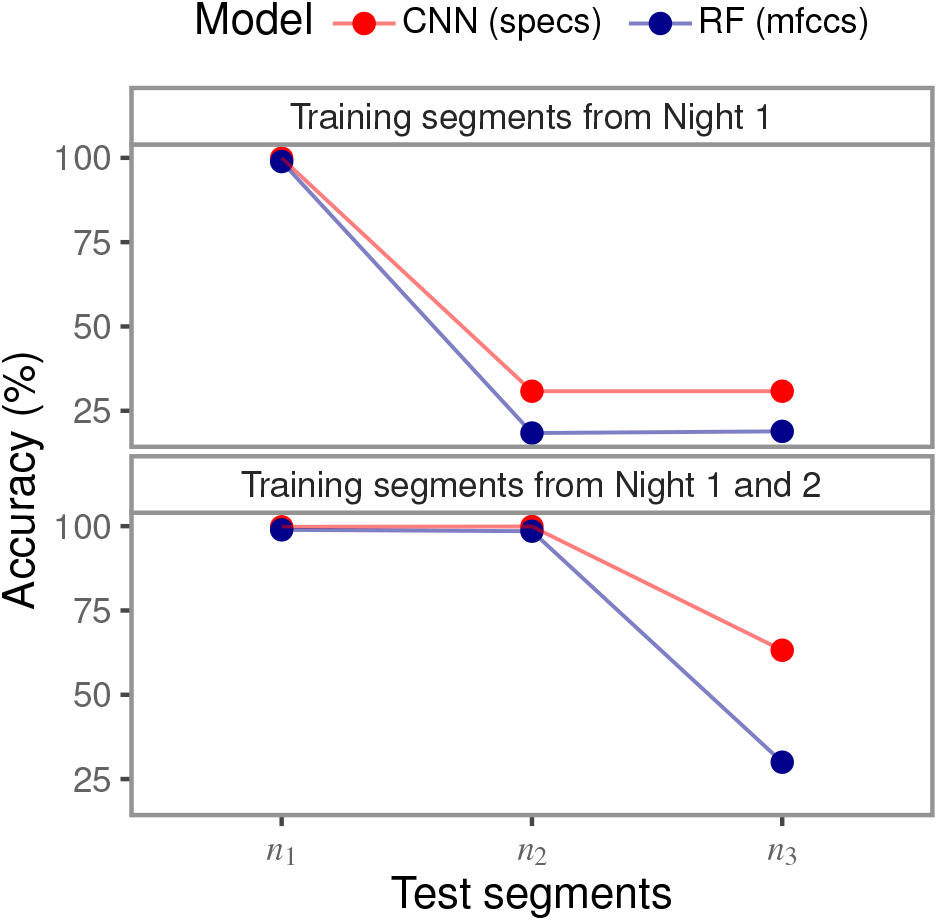
Assessment of model accuracy in recognizing cricket individuals within a closed population using 1s segments. *n*_1_, *n*_2_, and *n*_3_ denote samples from night 1, 2, and 3, respectively. Exact numerical results are provided in Supplementary Material Table D.1.

### 3.2 Open-population identification accuracy over new individuals

The best-performing models accurately identified cricket individuals from same-night and different-night pairs, with slightly better performance on same-night pairs. Because the results were very similar irrespective of whether test samples were from new individuals or not, and consistent across 1s segments and five-syllable chirps, we only present the results for 1s segments from new individuals, with the remainder covered in Supplementary Material A.2 and B.

CNN and RF models trained on segment pairs from the first night achieved high accuracy in identifying whether segment pairs from the same nights, (*n*_1_, *n*_1_), (*n*_2_, *n*_2_), and (*n*_3_, *n*_3_), belonged to the same individual, with average accuracies of 95.0% and 86.1%, respectively (Figure 4). Even when applied to segment pairs from different nights, (*n*_1_, *n*_2_), (*n*_1_, *n*_3_), and (*n*_2_, *n*_3_), the models still performed well, with average accuracies of 80.2% and 68.3%.

**Figure 4:**
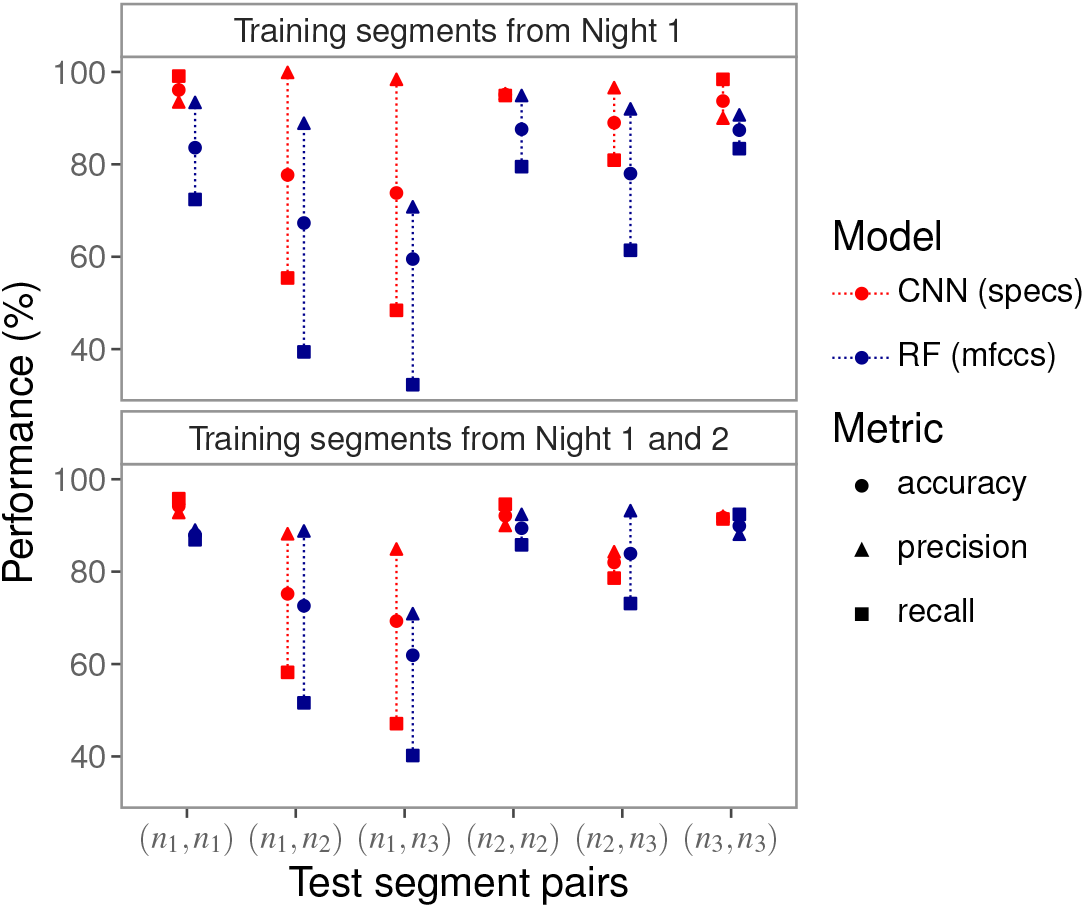
Assessment of model performance in identifying crickets within an open population using 1s segments from new individuals. Same-night pairs: (*n*_1_, *n*_1_), (*n*_2_, *n*_2_), and (*n*_3_, *n*_3_); different-night pairs: (*n*_1_, *n*_2_), (*n*_1_, *n*_3_), and (*n*_2_, *n*_3_). Exact numerical results are provided in Supplementary Material Table D.3.

Training on two-night segment data exclusively improved the performance of the RF models. Specifically, the RF model trained on segment pairs from the first and second nights increased the average identification accuracy by 3.0% on same-night pairs (night 1: 86.1%, nights 1 and 2: 89.1%) and 4.5% on different-night pairs (night 1: 68.3%, nights 1 and 2: 72.8%).

### 3.3 CNN vs RF models

CNNs outperformed RF models across all evaluated scenarios. In the closed-population identification task, CNNs demonstrated superior performance in all cases, with notable improvements observed when classifying individuals recorded on nights distinct from those used in training.

The CNN model trained on segments from night 1, increased the average recognition accuracy by 12.6% when classifying individuals from night 2 and 3 (CNN: 30.8%, RF: 18.6%). Similarly, the CNN model trained on segments from night 1 and 2, improved the recognition accuracy by 33.2% when classifying individuals from night 3 (CNN: 63.2%, RF: 30.0%).

For the open-population identification task, CNNs also outperformed RF in all cases. For example, the CNN model trained on segment pairs from night 1, increased the average identification accuracy by 8.9% on same-night pairs (CNN: 95.0%, RF: 86.1%) and 11.9% on different-night pairs (CNN: 80.2%, RF: 68.3%) than RF.

### 3.4 Temperature analysis

Increases in temperature were associated with shorter chirp duration (*β* = −133.6, s.e. = 1.4, *p* ≪ 0.001) and higher carrier frequencies (*β* = 66.6, s.e. = 0.7, *p* ≪ 0.001). Misclassification errors were positively associated with differences in temperature. When trained and tested on spectrograms from the same individuals with no adjustment for temperature, an increase in temperature of 0.5^*°*^*C* increased the odds of misclassification by 46.0% (95% CI 41.6–49.9%) if only night 1 recordings were used, and 13.8% (95% CI 7.7–20.2%) if recordings from night 1 and 2 were used. When trained and tested on spectrograms from different individuals, again without adjusting for temperature, the same temperature increase increased the odds of misclassification only if recordings from night 1 and 2 were used (by 46.0%, 95% CI 41.6–49.9%; night 1 only = 1.3%, 95% CI −4.3%–1.8%). Despite these apparent effects of temperature on misclassification error, adjusting spectrograms either for chirp duration or for both chirp duration and carrier frequency gave no improvement in performance, most likely due to substantial noise in the estimated relationship between temperature and acoustic features (Supplementary Material C).

## 4 Discussion

We used deep learning to evaluate the feasibility of recognizing individual crickets of the species *P. guttiventris* exclusively from their acoustic recordings in both closed and open population contexts. The closed-population context is one in which the identities of all individuals in the population are known *a priori*. The task in this case corresponds to a standard multi-class classification problem in which calls must be assigned to one of a fixed number of classes, each class representing an individual animal. In the open-population context, the number of individuals in the population is unknown and potentially changing over time. In this case, the task we addressed was the classification of pairs of calls as made by the same animal, or by different animals. These classifications can be used to identify individual animals by comparing calls to a reference catalog containing calls from known individuals (see e.g. Cheeseman et al. (2022) in the context of photographic identification).

In both closed- and open-population identification tasks, the best models demonstrated much higher accuracy in identifying cricket individuals from the same nights as training data than other nights (closed population: Figures 3 and A.1; open population: Figures 4 and A.2). For open-population models, this was true regardless of whether test pairs were drawn from new individuals (Figure 4 and Figure D.4) or individuals used in training (Figures D.5 and D.6). These findings support previous research by Nandi and Balakrishnan (2013) indicating that cricket intra-individual variability is lower within a night compared to across multiple nights.

One source of variability, and therefore source of misclassification error, is a change in temperature, which affects the duration and pitch of *P. guttiventris* chirps, with knock-on effects for spectrogram construction. Standard approaches for temperature normalization did not reduce misclassification errors, most likely because adjustments based on mean effects did not adequately capture individual heterogeneity. However, because temperature can be easily collected as part of passive monitoring, and more sophisticated temperature adjustments could possibly be learned as part of classifier training, this source of error may be reduced in future.

Training the model on a variety of samples from multiple nights enhances its robustness to intra-individual variability and consistently yields incremental improvements in identifying individual crickets across different nights, especially when training and test sets come from the same individuals (Figure 3, Figure A.1, Figure B.3, and Figure B.4). However, when training and test samples came from different individuals (Figure 4 and Figure A.2), an improvement was only observed for RF models. These findings suggest that with additional training recordings across more nights, classification accuracy in new, unseen nights would be higher than that reported here.

Although the literature documents many cricket studies exclusively relying on temporal features (Bertram, Fitzsimmons, McAuley, Rundle, & Gorelick, 2012; Ryder & Siva-Jothy, 2000), RF models using these features consistently achieved lower accuracy than models using spectrograms or MFCCs. Therefore, temporal features alone should not be used in isolation for this application, they should rather be combined with other features like MFCCs for improved accuracy as shown in Figure A.1.

Unlike standard classification models that cannot recognize previously unobserved individuals, similarity learning models demonstrated very similar accuracy for both new individuals (Figure 4 and Figure A.2) and those used in training (Figures B.3 and B.4). This confirms the ability of these models to generalize to previously unobserved individuals, making them suitable for open-population identification.

These results demonstrate both the feasibility of identifying crickets from their calls and the potential of deep learning as an effective and labor-efficient tool for this purpose. Although our models did not achieve perfect individual identification accuracy for fully automated use, they can still be integrated with manual processes in a semi-automated manner to accelerate annotation and maximize efficiency and accuracy. In the closed-population context, our deep learning classifiers can process fixed-length call segments or chirps and predict preliminary probabilities that samples belong to particular individuals. A predefined probability threshold can determine which predictions are reliable enough to be accepted automatically and which require human review. Predictions above the threshold can be accepted as they are, while those below the threshold can be subject to manual verification. This approach allows human experts to review, refine, and confirm the classifier’s predictions.

In the realm of open-population, similarity learning methods predict the similarity distance between pairs of samples, which can be useful in two practical scenarios. The first scenario involves assigning a new sample to an existing individual in the reference catalog or creating a new identity if it does not match any existing ones. In this context, the new sample is compared with representative samples from existing individuals in the catalog, and the pairs are ranked by their similarity distance. Excluding all pairs with a similarity distance above a predefined or optimized threshold as mismatches and only presenting the reduced subset of potential matches to the user for additional manual verification assists and speeds up manual individual identification. The second scenario entails identifying individual animals from a new set of samples with no existing reference catalog. In this framework, the first sample is compared with the remaining subsequent samples, and the produced pairs are again ranked by similarity distance. Once again, ruling out all pairs with a similarity distance exceeding a predetermined similarity threshold as mismatches before presenting the remaining potential matches to the user for manual confirmation supports and accelerates manual individual identification. Any confirmed matches would be documented, and the same process continues until all subsequent samples are exhausted. This second scenario is more challenging, and most existing practical applications and photo-matching software are typically limited to the first scenario.

Acoustic signals of insects and anurans are usually simple and repetitive in their temporal structure, with limited species-specific repertoires (Gerhardt & Huber, 2002). Among the orthopterans, field and tree cricket calls especially, have a narrow spectral bandwidth, with most of the energy concentrated around the carrier frequency. To our knowledge, this is the first study that has attempted AIID on an insect species with simple, less-featured/detailed signals. The high accuracy of our classifiers in identifying individual field crickets, firstly demonstrates the efficiency of these machine learning classifiers in classifying signals with a relatively limited set of features both in the spectral and temporal domains. Secondly, considering the structural similarity of acoustic signals across different cricket species, our classifiers can be used for investigating AIID in several orthopteran species. In fact, our deep learning based classifier is likely to perform better on katydid and grasshopper calls which have a broad spectral bandwidth as well as greater temporal complexity (Nityananda & Balakrishnan, 2006). Certain cicada and anuran species have also been reported to show larger species-specific acoustic repertoires (Gogala, 1995; Narins, Lewis, & McClelland, 2000), which can yield better AIID performance.

## 5 Conclusion

This study demonstrates the feasibility of noninvasively identifying cricket individuals of the species *P. guttiventris* from their calls in closed and open population contexts using deep learning methodologies. Using a three-step approach that integrates preprocessing, feature extraction, and identification, we successfully identified cricket individuals from same-night and different-night audio recordings using models trained on single and two-night data.

In closed population, the identification accuracy was higher when classifying individuals from the same nights as training data, while for closed population, it was higher for pairs drawn from the same nights. Amalgamating data from multiple nights increased model robustness to intra-individual variability, thus enhancing model ability in recognizing cricket individuals from different-night recordings. Deep learning outperformed RF models, but combining MFCCs and temporal features elevated the accuracy of RF models. The results demonstrate the feasibility of recognizing cricket individuals exclusively from their acoustic calls and the potential of deep learning models to support AIID for non-invasively monitoring insect species.

While our methods may not fully automate animal identification from calls, they can significantly assist and expedite manual identification within a semi-automatic framework. Integrating our classifiers with manual processes can enhance the scale and effectiveness of conservation efforts, enabling ecologists to focus more on addressing ecological issues.

## Data availability

The code for training and testing neural network and random forest models and a subset of the audio files are currently available at https://github.com/kabuga1987/Cricket_Identification/tree/main. Upon acceptance, we will use Zenodo to generate a permanent DOI link to the repository used to produce the manuscript’s results, and we will make the entire dataset publicly accessible.

## Acknowledgment

We thank the South African Centre for High performance Computing (CHPC) for providing us with computational resources. Emmanuel Kabuga is supported by a Doctoral Fellowship from the University of Cape Town.

