## Supplementary Material for "Acoustic individual identification in a species of field cricket using deep learning"

### A Individual identification accuracy from five-syllable chirps

#### A.1 Closed-population classification accuracy

The best convolutional neural network (CNN) and random forest (RF) models correctly classified chirps from the same nights they were trained on with nearly perfect accuracy, irrespective of whether they are trained on single- or two-night data (Figure A.1). When tested on different nights ($n_2$ and $n_3$), models trained on first-night chirps still outperformed random classifiers, with average accuracies of 31.6% (CNN) and 31.4% (RF).

Training on two-night chirps improved the classification accuracy of chirps from different nights as training data. For example, the CNN model increased the accuracy by 27.5% (night 1: 31.6%, nights 1 and 2: 59.1%) when classifying chirps from night 3, $n_3$.

Aggregating MFCCs and temporal features elevated the RF model performance, with training on MFCCs alone also yielding better results than using only temporal features (Figure A.1). For example, the RF model—using both MFCCs and temporal features—trained on two-night chirps increased the recognition accuracy over chirps from night 3 by 8.3% than MFCC-only model (RF (mfccs+tfs): 51.8%, RF (mfccs): 43.5%) and by 22.2% than the temporal feature-only model (RF(mfccs+tfs): 51.8%, RF(tfs): 29.6%).

Although the best CNN and RF models achieved comparable accuracy when classifying chirps from the same nights they were trained on, the CNN model trained on two-night data outperformed the best RF model by 7.3% in classifying chirps from night 3, $n_3$ (CNN: 59.1%, RF (mfccs + tfs): 51.8%).

#### A.2 Open-population identification accuracy

CNN and RF models trained on single-night or two-night chirp pairs achieved higher identification accuracy on same-night pairs—$(n_1, n_1)$, $(n_2, n_2)$, and $(n_3, n_3)$—than different-night pairs—$(n_1, n_2)$, $(n_1, n_3)$, and $(n_2, n_3)$ (Figure A.2). For example, the CNN model trained on single-night pairs increased the average identification accuracy by 15.3% for same-night pairs (94.1%) compared to different-night pairs (78.8%). Likewise, the RF model increased the identification accuracy by 20.3% over the above-mentioned pairs).

Training on two-night data improved the performance of the RF models only. Specifically, the RF model trained on two-night pairs increased average identification accuracy by 4.7% on same-night pairs (night 1: 85.4%, nights 1 and 2: 90.1%) and by 2.7% on different-night pairs (night 1: 65.2%, night 1 and 2: 67.9%).

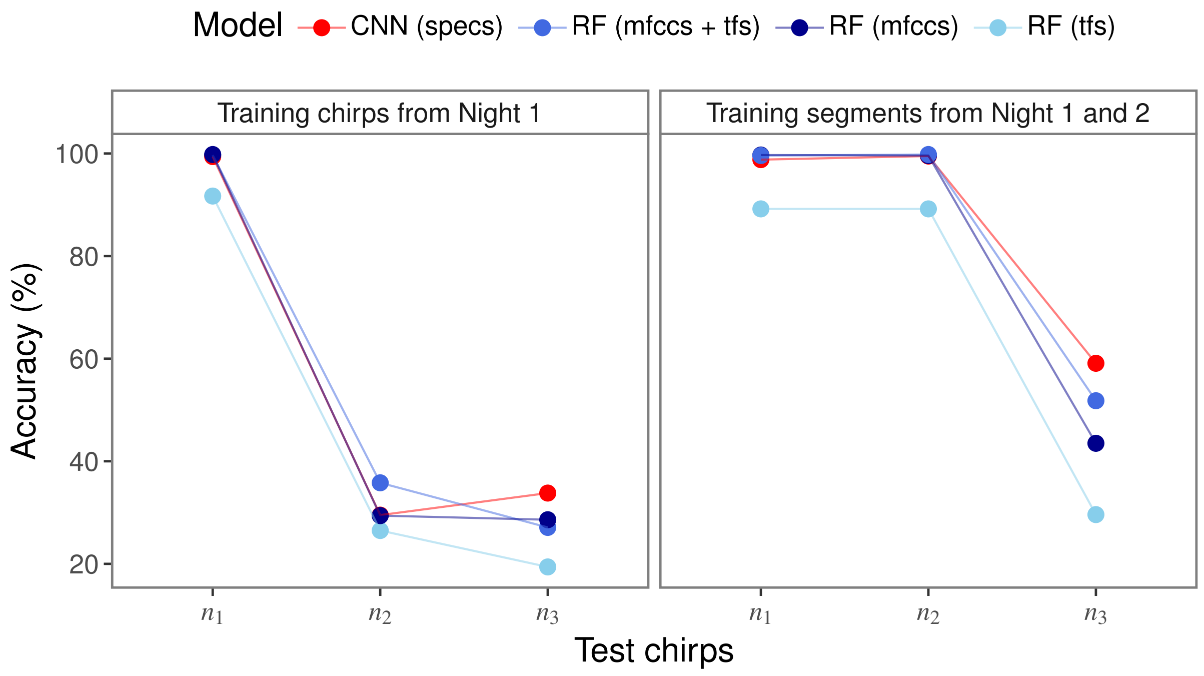


Figure A.1: Assessment of model accuracy in recognizing cricket individuals within a closed population using 5-syllable chirps. $n_1, n_2$ and $n_3$ denote test chirps from night 1, 2, and 3. Exact numeral results are provided in Supplementary Material Table D.2.

CNN models consistently outperformed RF models across all scenarios (Figure A.2). For instance, the CNN model trained on single-night pairs increased the identification accuracy by 8.7% for same-night pairs (CNN: 94.1%, RF: 84.5%) and 13.7% for different-night pairs (CNN: 78.8%, RF: 65.1%).

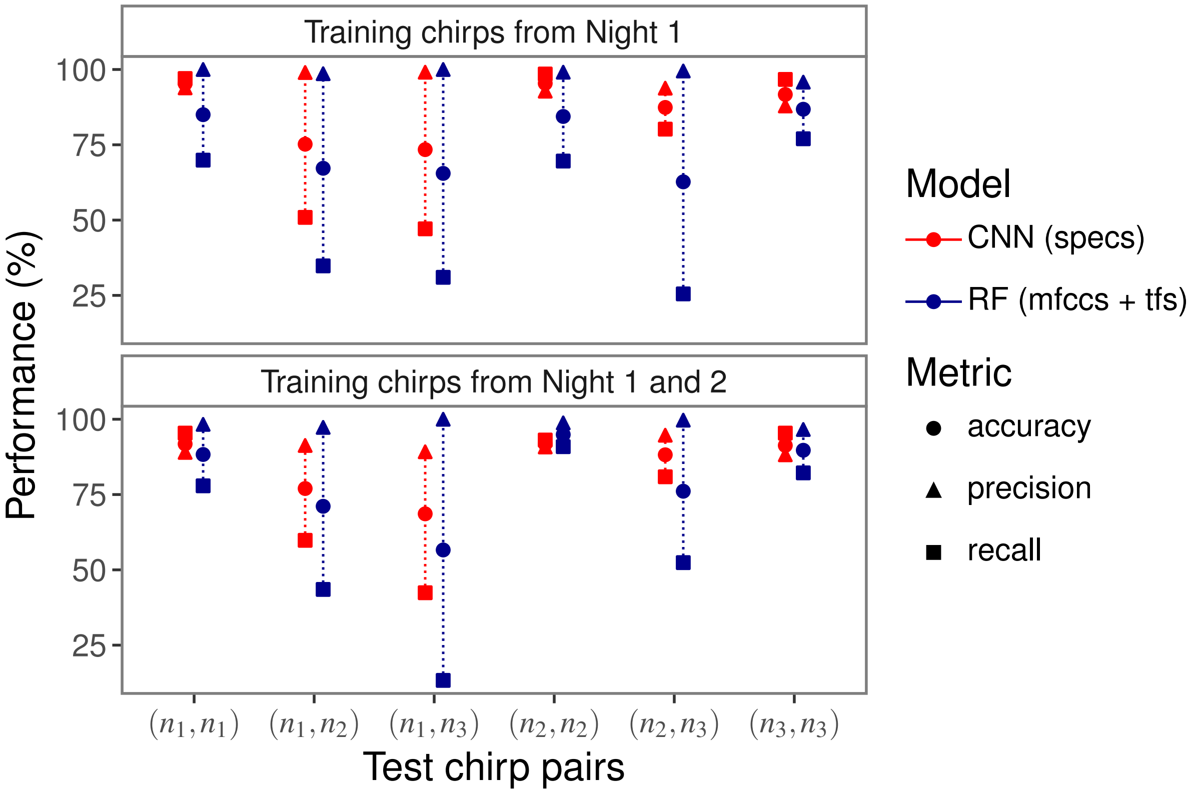


Figure A.2: Assessment of model performance in identifying crickets within an open population using 5-syllable segments from new individuals. Same-night pairs: $(n_1, n_1), (n_2, n_2)$, and $(n_3, n_3)$; different-night pairs: $(n_1, n_2), (n_1, n_3)$, and $(n_2, n_3)$. Exact numerical results are provided in Supplementary Material D.4.

### B Individual identification accuracy over known individuals

The best-performing models reliably identified individual crickets across same- and different-night pairs, with slightly higher accuracy on same-night pairs. Performance was consistent across 1s segments and five-syllable chirps, with CNNs outperforming RFs in 7 of 8 cases.

Models consistently achieved higher accuracy in identifying pairs from the same night compared to pairs from different nights, across both 1s segments and five-syllable chips, irrespective of whether models were trained on pairs from a single or two nights (Figures B.3 and B.4). For instance, CNN models trained on segments from night 1 correctly identified same-night pairs, $(n_1, n_1), (n_2, n_2)$ and $(n_3, n_3)$, with 95.1% average accuracy, while achieving 73% average accuracy for different-night pairs, $(n_1, n_2), (n_1, n_3)$ and $(n_2, n_3)$.

Training on two-night data (Figures B.3 and B.4) notably enhances different-night pair identification, improving average accuracy by 12.4–14.1% for 1s segments (CNN: 73% to 87.1%; RF: 67.2% to 79.6%) and 11.6–13.6% for five-syllable chirps (CNN: 72% to 83.6%; RF: 69% to 82.6%).

CNN models achieved higher accuracy than RF models, with a slightly higher margin on 1s segments than five-syllable chirps, more likely due to the inclusion of temporal and carrier frequency features for five-syllable chirp. For example, the CNN models trained on two-night 1s segments (Table B.3), increased the accuracy by 6.1–7.5% (same-night pairs: CNN (97%) vs. RF (90.9%), different-night pairs: CNN (87.1%) vs. RF (79.6%)) than RF models.

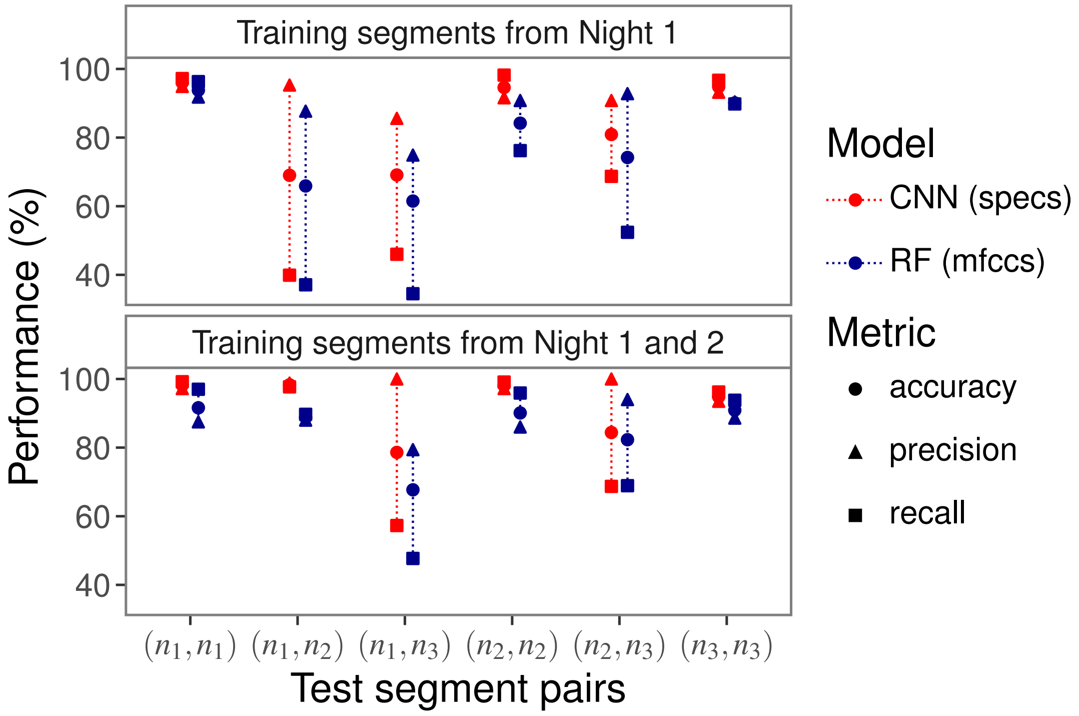


Figure B.3: Assessment of model performance in identifying crickets in an open population using 1s segments from test samples of the same individuals used in training. Exact numerical values are provided in Supplementary Material Table D.5. Same-night pairs: $(n_1, n_1), (n_2, n_2)$ and $(n_3, n_3)$; different-night pairs: $(n_1, n_2), (n_1, n_3)$ and $(n_2, n_3)$.

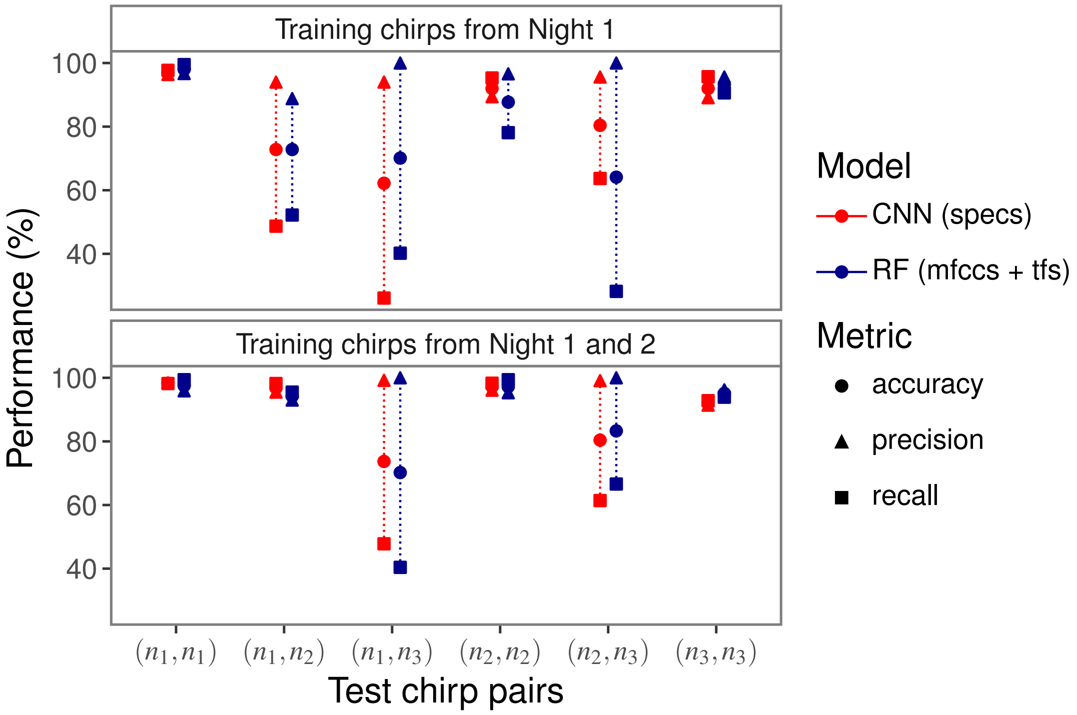


Figure B.4: Assessment of model performance in identifying crickets in an open population using five-syllable chirps from test samples of the same individuals used in training. Exact numerical values are provided in Supplementary Material Table D.6. Same-night pairs: $(n_1, n_1), (n_2, n_2)$ and $(n_3, n_3)$; different-night pairs: $(n_1, n_2), (n_1, n_3)$ and $(n_2, n_3)$.

### C Comparison of CNN performance on raw, duration-corrected, and duration- and frequency-corrected chirps

Our analysis showed that high temperature is associated with shorter chirp durations and higher carrier frequencies (Figure C.5). However, despite these apparent relationships, adjusting the spectrograms for chirp duration alone—or for both chirp duration and frequency—did not improve the performance of the best CNN model in either closed- or open-population settings. In most cases, identification accuracy was slightly higher when using raw chirp data compared to duration- and frequency-adjusted chirps (Figures C.6, C.7, and C.8).

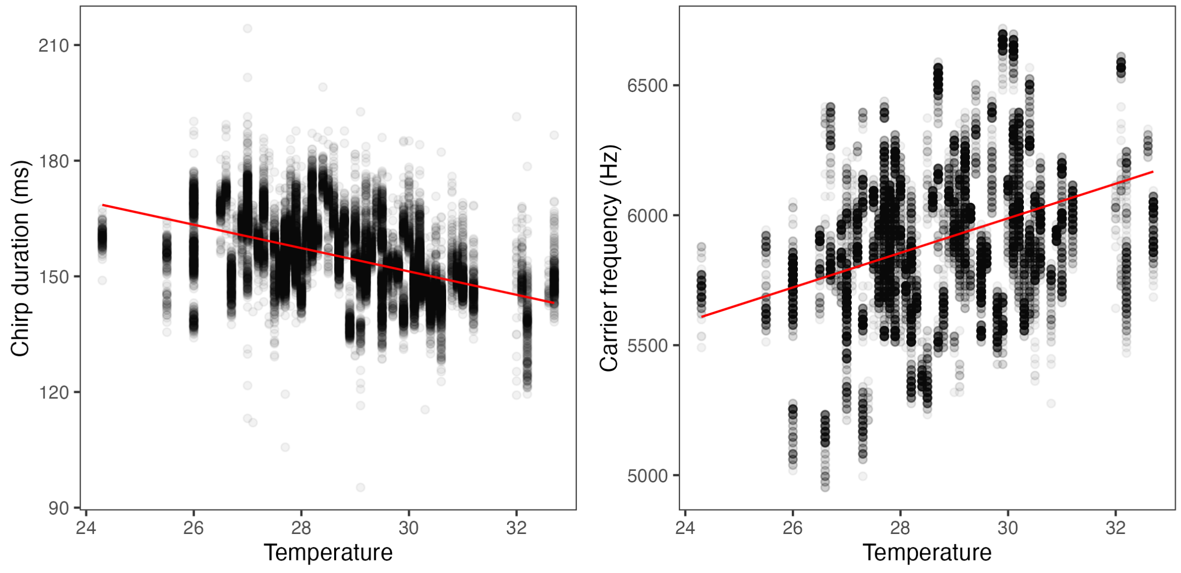


Figure C.5: Increases in temperature are associated with shorter chirp durations and higher carrier frequencies. Black circles show individual data points, with red lines showing lines of best fit from estimated GLMMs. The substantial variability in responses at any given temperature makes any temperature adjustments based on mean relationships unlikely to substantially improve classifier performance.

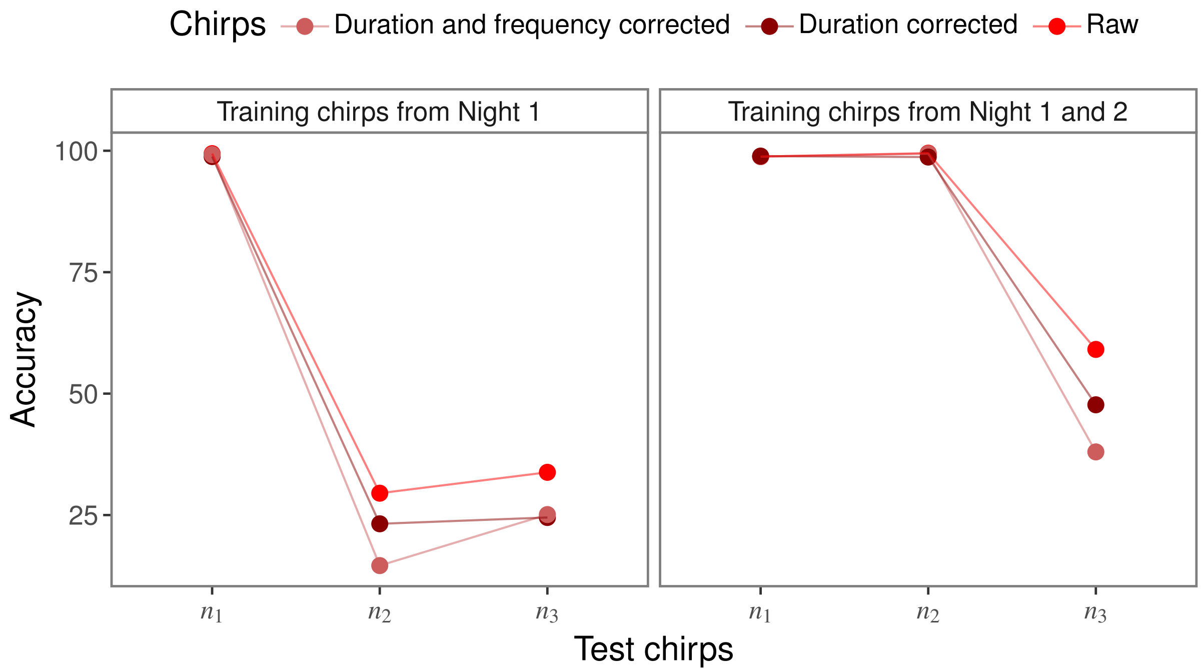


Figure C.6: Accuracy of best CNN models in classifying cricket individuals from raw, duration-corrected, and duration- and frequency-corrected five-syllable chirps within a closed population. $n_1, n_2$ and $n_3$ denote samples from night 1, 2, and 3. Exact numerical results are provided in Supplementary Material D.7.

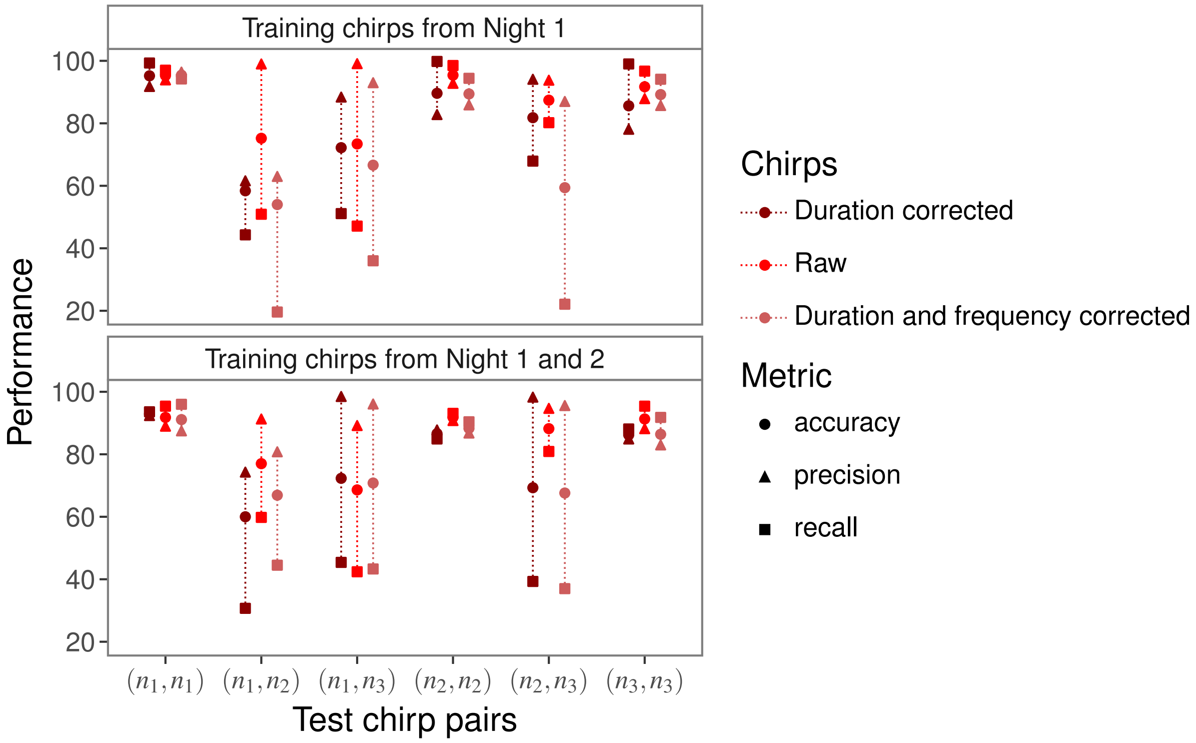


Figure C.7: Assessment of the performance of best CNN models in identifying cricket individuals from raw, duration-corrected, and duration- and frequency-corrected five-syllable chirps of new individuals within an open population. Same-night pairs: $(n_1, n_1), (n_2, n_3)$ and $(n_3, n_3)$; different-night pairs: $(n_1, n_2), (n_1, n_{,3})$ and $(n_2, n_3)$. Exact numerical results are provided in Supplementary Material D.8.

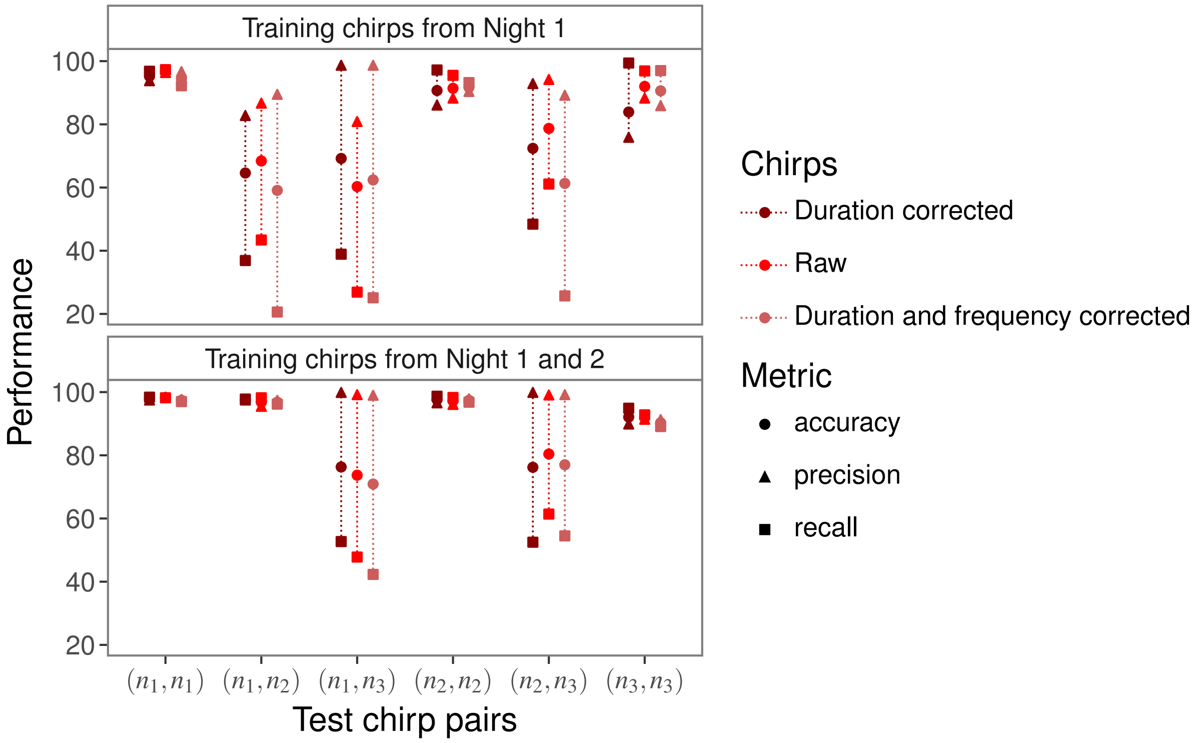


Figure C.8: Assessment of the performance of best CNN models in identifying cricket individuals from raw, duration-corrected, and duration- and frequency-corrected five-syllable chirps of known individuals within an open population. Same-night pairs: $(n_1, n_1), (n_2, n_3)$ and $(n_3, n_3)$; different-night pairs: $(n_1, n_2), (n_1, n_3)$ and $(n_2, n_3)$. Exact numerical results are provided in Supplementary Material Table D.9.

### D Exact numerical results

For completeness, and to address the difficulty of extracting precise numerical values from figures, we provide tabulated equivalents for all figures presented in both the main manuscript and the supplementary materials. Specifically, Tables D1 through D9 contain the exact numerical values corresponding to the following figures:

- Table D.1 : Figure 3
- Table D.2 : Figure A.1
- Table D.3 : Figure 4
- Table D.4 : Figure A.2
- Table D.5 : Figure B.3
- Table D.6 : Figure B.4
- Table D.7 : Figure C.6
- Table D.8 : Figure C.7
- Table D.9 : Figure C.8

| Models | Train | Test | Accuracy | Average accuracy |
|---|---|---|---|---|
| Single-night models | | | | |
| CNN (specs) | $n_1^{tr}$ | $n_1^{te}$ | 99.9 | **99.9** |
| | | $n_2^{te}$ | 30.8 | **30.8** |
| | | $n_3^{te}$ | 30.8 | |
| RF (mfccs) | $n_1^{tr}$ | $n_1^{te}$ | 98.9 | 98.9 |
| | | $n_2^{te}$ | 18.4 | 18.6 |
| | | $n_3^{te}$ | 18.9 | |
| Two-night models | | | | |
| CNN (specs) | $n_1^{tr}$ $n_2^{tr}$ | $n_1^{te}$ | 99.8 | **99.8** |
| | | $n_2^{te}$ | 99.9 | |
| | | $n_3^{te}$ | 63.2 | **63.2** |
| RF (mfccs) | $n_1^{tr}$ $n_2^{tr}$ | $n_1^{te}$ | 98.9 | 98.7 |
| | | $n_2^{te}$ | 98.5 | |
| | | $n_3^{te}$ | 30.0 | 30.0 |

Table D.1: Assessment of model performance in recognizing cricket individuals within a closed population using 1s segments.

| Models | Train | Test | Accuracy | Average accuracy |
| --- | --- | --- | --- | --- |
| Single-night models | | | | |
| CNN (specs) | $n_1^{tr}$ | $n_1^{te}$ | 99.4 | 99.4 |
| | | $n_2^{te}$ | 29.5 | **31.6** |
| | | $n_3^{te}$ | 33.8 | |
| RF (tfs) | $n_1^{tr}$ | $n_1^{te}$ | 91.7 | 91.7 |
| | | $n_2^{te}$ | 26.5 | 23.0 |
| | | $n_3^{te}$ | 19.4 | |
| RF (mfccs) | $n_1^{tr}$ | $n_1^{te}$ | 99.8 | **99.8** |
| | | $n_2^{te}$ | 29.4 | 29 |
| | | $n_3^{te}$ | 28.6 | |
| RF (mfccs + tfs) | $n_1^{tr}$ | $n_1^{te}$ | 99.7 | 99.7 |
| | | $n_2^{te}$ | 35.8 | 31.4 |
| | | $n_{3s}^{te}$ | 27.1 | |
| Two-night models | | | | |
| CNN (specs) | $n_1^{tr}$, $n_2^{tr}$ | $n_1^{te}$ | 98.8 | 99.2 |
| | | $n_2^{te}$ | 99.5 | |
| | | $n_3^{te}$ | 59.1 | **59.1** |
| RF (tfs) | $n_1^{tr}$, $n_2^{tr}$ | $n_1^{te}$ | 89.2 | 89.2 |
| | | $n_2^{te}$ | 89.2 | |
| | | $n_3^{te}$ | 29.6 | 29.6 |
| RF (mfccs) | $n_1^{tr}$, $n_2^{tr}$ | $n_1^{te}$ | 99.7 | **99.7** |
| | | $n_2^{te}$ | 99.6 | |
| | | $n_3^{te}$ | 43.5 | 43.5 |
| RF (mfccs + tfs) | $n_1^{tr}$, $n_2^{tr}$ | $n_1^{te}$ | 99.6 | **99.7** |
| | | $n_2^{te}$ | 99.8 | |
| | | $n_3^{te}$ | 51.8 | 51.8 |

Table D.2: Assessment of model performance in recognizing individual crickets within a closed population using five-syllable chirps.

| Models | Train | Test | Precision | Recall | F1 score | Accuracy | Average accuracy |
|---|---|---|---|---|---|---|---|
| | | | Single-night models | | | | |
| | | $(n_1^{te}, n_1^{te})$ | 93.5 | 99.1 | 96.2 | 96.1 | |
| | | $(n_2^{te}, n_2^{te})$ | 95.3 | 94.9 | 95.1 | 95.1 | **95.0** |
| CNN (specs) | | $(n_3^{te}, n_3^{te})$ | 90.0 | 98.4 | 94.0 | 93.7 | |
| | $(n_1^{tr}, n_1^{tr})$ | $(n_1^{te}, n_2^{te})$ | 99.9 | 55.4 | 71.3 | 77.7 | |
| | | $(n_1^{te}, n_3^{te})$ | 98.4 | 48.4 | 64.9 | 73.8 | **80.2** |
| | | $(n_2^{te}, n_3^{te})$ | 96.6 | 80.9 | 88.1 | 89.0 | |
| | | $(n_1^{te}, n_3^{te})$ | 93.4 | 72.4 | 81.6 | 83.6 | |
| | | $(n_2^{te}, n_2^{te})$ | 94.9 | 79.5 | 86.6 | 87.6 | 86.1 |
| RF (mfccs) | | $(n_3^{te}, n_3^{te})$ | 90.7 | 83.4 | 86.9 | 87.4 | |
| | $(n_1^{tr}, n_1^{tr})$ | $(n_1^{te}, n_2^{te})$ | 88.9 | 39.4 | 54.6 | 67.3 | |
| | | $(n_1^{te}, n_3^{te})$ | 70.8 | 32.3 | 44.3 | 59.5 | 68.3 |
| | | $(n_2^{te}, n_3^{te})$ | 92.0 | 61.4 | 73.6 | 78.0 | |
| | | | Two-night models | | | | |
| | $(n_1^{tr}, n_1^{tr})$ | $(n_1^{te}, n_1^{te})$ | 92.8 | 95.8 | 94.3 | 94.2 | |
| | | $(n_2^{te}, n_2^{te})$ | 90.0 | 94.6 | 92.3 | 92.2 | **92.7** |
| CNN (specs) | | $(n_3^{te}, n_3^{te})$ | 92.0 | 91.4 | 91.7 | 91.7 | |
| | $(n_1^{tr}, n_2^{tr})$ | $(n_1^{te}, n_2^{te})$ | 88.2 | 58.2 | 70.2 | 75.2 | |
| | | $(n_1^{te}, n_3^{te})$ | 84.9 | 47.1 | 60.6 | 69.3 | **75.5** |
| | $(n_2^{tr}, n_2^{tr})$ | $(n_2^{te}, n_3^{te})$ | 84.3 | 78.6 | 81.4 | 82.0 | |
| | $(n_1^{tr}, n_1^{tr})$ | $(n_1^{te}, n_1^{te})$ | 89.0 | 86.9 | 87.9 | 88.0 | |
| | | $(n_2^{te}, n_2^{te})$ | 92.4 | 85.8 | 89.0 | 89.4 | 89.1 |
| RF (mfccs + tfs) | | $(n_3^{te}, n_3^{te})$ | 88.1 | 92.4 | 90.1 | 89.9 | |
| | $(n_1^{tr}, n_2^{tr})$ | $(n_1^{te}, n_2^{te})$ | 88.8 | 51.6 | 65.3 | 72.6 | |
| | | $(n_1^{te}, n_3^{te})$ | 70.9 | 40.2 | 51.3 | 61.9 | 72.8 |
| | $(n_2^{tr}, n_2^{tr})$ | $(n_2^{te}, n_3^{te})$ | 93.2 | 73.1 | 81.9 | 83.9 | |

Table D.3: Assessment of model performance in identifying crickets within an open population using 1s segments from new individuals.

| Models | Train | Test | Precision | Recall | F1 score | Accuracy | Average accuracy |
|---|---|---|---|---|---|---|---|
| Single-night models | | | | | | | |
| CNN (specs) | $(n_1^{tr}, n_1^{tr})$ | $(n_1^{te}, n_1^{te})$ | 93.9 | 97.0 | 95.4 | 95.3 | **94.1** |
| | | $(n_2^{te}, n_2^{te})$ | 92.8 | 98.5 | 95.6 | 95.4 | |
| | | $(n_3^{te}, n_3^{te})$ | 87.9 | 96.7 | 92.1 | 91.7 | |
| | | $(n_1^{te}, n_2^{te})$ | 99.0 | 50.9 | 67.2 | 75.2 | **78.8** |
| | | $(n_1^{te}, n_3^{te})$ | 99.1 | 47.1 | 63.9 | 73.7 | |
| | | $(n_2^{te}, n_3^{te})$ | 93.8 | 80.2 | 86.5 | 87.4 | |
| RF (mfccs + tfs) | $(n_1^{tr}, n_1^{tr})$ | $(n_1^{te}, n_1^{te})$ | 100.0 | 69.9 | 82.3 | 85.0 | 85.4 |
| | | $(n_2^{te}, n_2^{te})$ | 99.1 | 69.6 | 81.7 | 84.4 | |
| | | $(n_3^{te}, n_3^{te})$ | 95.8 | 77.0 | 85.3 | 86.8 | |
| | | $(n_1^{te}, n_2^{te})$ | 98.6 | 34.8 | 51.4 | 67.2 | 65.1 |
| | | $(n_1^{te}, n_3^{te})$ | 100.0 | 31.0 | 47.3 | 65.5 | |
| | | $(n_2^{te}, n_3^{te})$ | 95.5 | 25.2 | 40.6 | 62.7 | |
| Two-night models | | | | | | | |
| CNN (specs) | $(n_1^{tr}, n_1^{tr})$ | $(n_1^{te}, n_1^{te})$ | 89.0 | 95.4 | 92.1 | 91.8 | **91.6** |
| | | $(n_2^{te}, n_2^{te})$ | 90.8 | 93.1 | 91.9 | 91.8 | |
| | $(n_1^{tr}, n_2^{tr})$ | $(n_3^{te}, n_3^{te})$ | 88.2 | 95.4 | 91.7 | 91.3 | |
| | | $(n_1^{te}, n_2^{te})$ | 91.3 | 59.8 | 72.2 | 77.0 | **77.9** |
| | | $(n_1^{te}, n_3^{te})$ | 89.2 | 42.4 | 57.5 | 68.6 | |
| | $(n_2^{tr}, n_2^{tr})$ | $(n_2^{te}, n_3^{te})$ | 94.7 | 80.9 | 87.2 | 88.2 | |
| RF (mfccs + tfs) | $(n_1^{tr}, n_1^{tr})$ | $(n_1^{te}, n_1^{te})$ | 98.3 | 77.9 | 86.9 | 88.3 | 90.1 |
| | | $(n_2^{te}, n_2^{te})$ | 98.8 | 90.9 | 94.7 | 94.9 | |
| | $(n_1^{tr}, n_2^{tr})$ | $(n_3^{te}, n_3^{te})$ | 96.6 | 82.2 | 88.8 | 89.7 | |
| | | $(n_1^{te}, n_2^{te})$ | 97.3 | 43.5 | 60.1 | 71.1 | 67.9 |
| | | $(n_1^{te}, n_3^{te})$ | 100.0 | 13.3 | 23.5 | 56.6 | |
| | $(n_2^{tr}, n_2^{tr})$ | $(n_2^{te}, n_3^{te})$ | 99.7 | 52.4 | 68.7 | 76.1 | |

Table D.4: Assessment of model performance in identifying crickets within an open population using five-syllable chirps from new individuals.

| Models | Train | Test | Precision | Recall | F1 score | Accuracy | Average accuracy |
|---|---|---|---|---|---|---|---|
| (a)Single-night models | | | | | | | |
| | | $(n_1^{te}, n_1^{te})$ | 94.9 | 97.2 | 96.0 | 96.0 | |
| | | $(n_2^{te}, n_2^{te})$ | 91.6 | 98.2 | 94.8 | 94.6 | **95.1** |
| CNN (specs) | $(n_1^{tr}, n_1^{tr})$ | $(n_3^{te}, n_3^{te})$ | 93.2 | 96.7 | 94.9 | 94.8 | |
| | | $(n_1^{te}, n_2^{te})$ | 95.3 | 39.8 | 56.3 | 69.0 | |
| | | $(n_1^{te}, n_3^{te})$ | 85.6 | 46.0 | 59.8 | 69.1 | **73** |
| | | $(n_2^{te}, n_3^{te})$ | 90.8 | 68.7 | 78.2 | 80.9 | |
| | | $(n_1^{te}, n_1^{te})$ | 91.8 | 96.3 | 94.0 | 93.8 | |
| | | $(n_2^{te}, n_2^{te})$ | 90.8 | 76.2 | 82.8 | 84.2 | 89.4 |
| RF (mfccs + tfs) | $(n_1^{tr}, n_1^{tr})$ | $(n_3^{te}, n_3^{te})$ | 90.4 | 89.8 | 90.1 | 90.1 | |
| | | $(n_1^{te}, n_2^{te})$ | 87.7 | 37.1 | 52.1 | 65.9 | |
| | | $(n_1^{te}, n_3^{te})$ | 74.9 | 34.5 | 47.2 | 61.5 | 67.2 |
| | | $(n_2^{te}, n_3^{te})$ | 92.8 | 52.4 | 67.0 | 74.2 | |
| (b)Two-night models | | | | | | | |
| | $(n_1^{tr}, n_1^{tr})$ | $(n_1^{te}, n_1^{te})$ | 97.2 | 99.2 | 98.2 | 98.2 | |
| | | $(n_2^{te}, n_2^{te})$ | 97.2 | 99.1 | 98.2 | 98.1 | **97.0** |
| CNN (specs) | $(n_1^{tr}, n_2^{tr})$ | $(n_3^{te}, n_3^{te})$ | 93.5 | 96.2 | 94.8 | 94.8 | |
| | | $(n_1^{te}, n_2^{te})$ | 98.8 | 97.7 | 98.3 | 98.3 | |
| | | $(n_1^{te}, n_3^{te})$ | 100.0 | 57.3 | 72.9 | 78.6 | **87.1** |
| | $(n_2^{tr}, n_2^{tr})$ | $(n_2^{te}, n_3^{te})$ | 100.0 | 68.7 | 81.5 | 84.4 | |
| | $(n_1^{tr}, n_1^{tr})$ | $(n_1^{te}, n_{1s}^{te})$ | 87.5 | 97.0 | 92.0 | 91.6 | |
| | | $(n_2^{te}, n_2^{te})$ | 86.0 | 95.9 | 90.7 | 90.1 | 90.9 |
| RF (mfccs + tfs) | $(n_1^{tr}, n_2^{tr})$ | $(n_3^{te}, n_3^{te})$ | 88.6 | 93.8 | 91.1 | 90.9 | |
| | | $(n_1^{te}, n_2^{te})$ | 88.0 | 89.7 | 88.9 | 88.8 | |
| | | $(n_1^{te}, n_3^{te})$ | 79.4 | 47.7 | 59.6 | 67.7 | 79.6 |
| | $(n_2^{tr}, n_2^{tr})$ | $(n_2^{te}, n_3^{te})$ | 94.0 | 68.9 | 79.5 | 82.3 | |

Table D.5: Assessment of model performance in identifying crickets in a open population using 1s segments from test samples of the same individuals used in training.

| Models | Train | Test | Precision | Recall | F1 score | Accuracy | Average accuracy |
|---|---|---|---|---|---|---|---|
| | | | (a)Single-night models | | | | |
| CNN (specs) | $(n_1^{tr}, n_1^{tr})$ | $(n_1^{te}, n_1^{te})$ | 96.4 | 97.7 | 97.0 | 97.0 | |
| | | $(n_2^{te}, n_2^{te})$ | 89.4 | 95.3 | 92.3 | 92.0 | **93.7** |
| | | $(n_3^{te}, n_3^{te})$ | 89.1 | 95.7 | 92.3 | 92.0 | |
| | | $(n_1^{te}, n_2^{te})$ | 94.0 | 48.7 | 64.1 | 72.8 | |
| | | $(n_1^{te}, n_3^{te})$ | 94.0 | 26.1 | 40.8 | 62.8 | **72** |
| | | $(n_2^{te}, n_3^{te})$ | 95.6 | 63.7 | 76.4 | 80.4 | |
| RF (mfccs + tfs) | $(n_1^{tr}, n_1^{tr})$ | $(n_1^{te}, n_1^{te})$ | 96.7 | 99.5 | 98.1 | 98.0 | |
| | | $(n_2^{te}, n_2^{te})$ | 96.6 | 78.1 | 86.4 | 87.7 | 93.0 |
| | | $(n_3^{te}, n_3^{te})$ | 95.6 | 90.6 | 93.0 | 93.2 | |
| | | $(n_1^{te}, n_2^{te})$ | 88.8 | 52.2 | 65.8 | 72.8 | |
| | | $(n_1^{te}, n_3^{te})$ | 100.0 | 40.2 | 57.4 | 70.1 | 69.0 |
| | | $(n_2^{te}, n_3^{te})$ | 1100.0 | 28.2 | 44.0 | 64.1 | |
| | | | (b)Two-night models | | | | |
| CNN (specs) | $(n_1^{tr}, n_1^{tr})$ | $(n_1^{te}, n_1^{te})$ | 98.4 | 98.2 | 98.3 | 98.3 | |
| | | $(n_2^{te}, n_2^{te})$ | 96.1 | 98.3 | 97.2 | 97.2 | 95.8 |
| | $(n_1^{tr}, n_2^{tr})$ | $(n_3^{te}, n_3^{te})$ | 91.4 | 92.8 | 92.0 | 92.0 | |
| | | $(n_1^{te}, n_2^{te})$ | 95.5 | 98.2 | 96.8 | 96.8 | |
| | | $(n_1^{te}, n_3^{te})$ | 99.2 | 47.8 | 64.5 | 73.7 | **83.6** |
| | $(n_2^{tr}, n_2^{tr})$ | $(n_2^{te}, n_3^{te})$ | 99.1 | 61.4 | 75.8 | 80.4 | |
| RF (mfccs + tfs) | $(n_1^{tr}, n_1^{tr})$ | $(n_1^{te}, n_1^{te})$ | 95.9 | 99.4 | 97.6 | 97.5 | |
| | | $(n_2^{te}, n_2^{te})$ | 95.3 | 99.4 | 97.3 | 97.2 | **96.6** |
| | $(n_1^{tr}, n_2^{tr})$ | $(n_3^{te}, n_3^{te})$ | 96.2 | 93.9 | 95.0 | 95.1 | |
| | | $(n_1^{te}, n_2^{te})$ | 93.0 | 95.5 | 94.2 | 94.2 | |
| | | $(n_1^{te}, n_3^{te})$ | 100.0 | 40.4 | 57.6 | 70.2 | 82.6 |
| | $(n_2^{tr}, n_2^{tr})$ | $(n_2^{te}, n_3^{te})$ | 100.0 | 66.6 | 80.0 | 83.3 | |

Table D.6: Assessment of model performance in identifying cricket individuals in an open population using five-syllable chirps from test samples of the same individuals used in training.

| Data | Train | Test | Accuracy | Average accuracy |
|---|---|---|---|---|
| Single-night models | | | | |
| Raw | $n_1^{tr}$ | $n_1^{te}$ | 99.4 | **99.4** |
| | | $n_2^{te}$ | 29.5 | **31.6** |
| | | $n_3^{te}$ | 33.8 | |
| Duration corrected | $n_1^{tr}$ | $n_1^{te}$ | 98.8 | 98.8 |
| | | $n_2^{te}$ | 23.2 | 23.9 |
| | | $n_3^{te}$ | 24.5 | |
| Duration and frequency corrected | $n_1^{tr}$ | $n_1^{te}$ | 99.2 | 99.2 |
| | | $n_2^{te}$ | 14.6 | 19.9 |
| | | $n_3^{te}$ | 25.1 | |
| Two-night models | | | | |
| Raw | $n_1^{tr}$ | $n_1^{te}$ | 98.8 | **99.2** |
| | $n_2^{tr}$ | $n_2^{te}$ | 99.5 | |
| | | $n_3^{te}$ | 59.1 | **59.1** |
| Duration corrected | $n_1^{tr}$ | $n_1^{te}$ | 98.9 | 98.8 |
| | $n_2^{tr}$ | $n_2^{te}$ | 98.7 | |
| | | $n_3^{te}$ | 47.7 | 47.7 |
| Duration and frequency corrected | $n_1^{tr}$ | $n_1^{te}$ | 98.8 | 99.1 |
| | $n_2^{tr}$ | $n_2^{te}$ | 99.4 | |
| | | $n_3^{te}$ | 38.0 | 38.0 |

Table D.7: Accuracy of best CNN models in classifying cricket individuals from raw, duration-corrected, and duration- and frequency-corrected five-syllable chirps within a closed population.

| Data | Train | Test | Precision | Recall | F1 score | Accuracy | Average accuracy |
|---|---|---|---|---|---|---|---|
| Single-night models | | | | | | | |
| Raw | $(n_1^{tr}, n_1^{tr})$ | $(n_1^{te}, n_1^{te})$ | 93.9 | 97.0 | 95.4 | 95.3 | |
| | | $(n_2^{te}, n_2^{te})$ | 92.8 | 98.5 | 95.6 | 95.4 | **94.1** |
| | | $(n_3^{te}, n_3^{te})$ | 87.9 | 96.7 | 92.1 | 91.7 | |
| | | $(n_1^{te}, n_2^{te})$ | 99.0 | 50.9 | 67.2 | 75.2 | |
| | | $(n_1^{te}, n_3^{te})$ | 99.1 | 47.1 | 63.9 | 73.4 | **78.7** |
| | | $(n_2^{te}, n_3^{te})$ | 93.8 | 80.2 | 86.5 | 87.4 | |
| Duration corrected | $(n_1^{tr}, n_1^{tr})$ | $(n_1^{te}, n_1^{te})$ | 91.0 | 99.3 | 95.4 | 95.2 | |
| | | $(n_2^{te}, n_2^{te})$ | 82.8 | 99.8 | 90.5 | 89.6 | 90.1 |
| | | $(n_3^{te}, n_3^{te})$ | 78.1 | 99.0 | 87.3 | 85.6 | |
| | | $(n_1^{te}, n_2^{te})$ | 61.6 | 44.3 | 51.5 | 58.4 | |
| | | $(n_1^{te}, n_3^{te})$ | 88.4 | 51.1 | 64.8 | 72.2 | 70.8 |
| | | $(n_2^{te}, n_3^{te})$ | 94.1 | 67.9 | 78.9 | 81.8 | |
| Duration and frequency corrected | $(n_1^{tr}, n_1^{tr})$ | $(n_1^{te}, n_1^{te})$ | 96.4 | 94.2 | 95.3 | 95.4 | |
| | | $(n_2^{te}, n_2^{te})$ | 85.9 | 94.4 | 90. | 89.4 | 91.3 |
| | | $(n_3^{te}, n_3^{te})$ | 85.7 | 94.1 | 89.7 | 89.2 | |
| | | $(n_1^{te}, n_2^{te})$ | 63.0 | 19.6 | 29.9 | 54.0 | |
| | | $(n_1^{te}, n_3^{te})$ | 93.0 | 36.0 | 51.9 | 66.6 | 60 |
| | | $(n_2^{te}, n_3^{te})$ | 87.0 | 22.1 | 35.3 | 59.4 | |
| Two-night models | | | | | | | |
| Raw | $(n_1^{tr}, n_1^{tr})$ | $(n_1^{te}, n_1^{te})$ | 89.0 | 95.4 | 92.1 | 91.8 | |
| | | $(n_2^{te}, n_2^{te})$ | 90.8 | 93.1 | 91.9 | 91.8 | **91.6** |
| | $(n_1^{tr}, n_2^{tr})$ | $(n_3^{te}, n_3^{te})$ | 88.2 | 95.4 | 91.7 | 91.3 | |
| | | $(n_1^{te}, n_2^{te})$ | 91.3 | 59.8 | 72.2 | 77.0 | |
| | | $(n_1^{te}, n_3^{te})$ | 89.2 | 42.4 | 57.5 | 68.6 | **77.9** |
| | $(n_2^{tr}, n_2^{tr})$ | $(n_2^{te}, n_3^{te})$ | 94.7 | 80.9 | 87.2 | 88.2 | |
| Duration corrected | $(n_1^{tr}, n_1^{tr})$ | $(n_1^{te}, n_1^{te})$ | 92.4 | 93.6 | 93.0 | 92.9 | |
| | | $(n_2^{te}, n_2^{te})$ | 87.8 | 84.9 | 86.3 | 86.6 | 86.6 |
| | $(n_1^{tr}, n_2^{tr})$ | $(n_3^{te}, n_3^{te})$ | 84.9 | 88.1 | 86.5 | 86.2 | |
| | | $(n_1^{te}, n_2^{te})$ | 74.3 | 30.7 | 43.4 | 60.0 | |
| | | $(n_1^{te}, n_3^{te})$ | 98.5 | 45.4 | 62.1 | 72.3 | 67.2 |
| | $(n_2^{tr}, n_2^{tr})$ | $(n_2^{te}, n_3^{te})$ | 98.3 | 39.3 | 56.2 | 69.3 | |
| Duration and frequency corrected | $(n_1^{tr}, n_1^{tr})$ | $(n_1^{te}, n_1^{te})$ | 87.5 | 96.0 | 91.6 | 91.1 | |
| | | $(n_2^{te}, n_2^{te})$ | 86.8 | 90.4 | 88.6 | 88.3 | 88.6 |
| | $(n_1^{tr}, n_2^{tr})$ | $(n_3^{te}, n_3^{te})$ | 83.0 | 91.8 | 87.1 | 86.4 | |
| | | $(n_1^{te}, n_2^{te})$ | 80.8 | 44.5 | 57.4 | 66.9 | |
| | | $(n_1^{te}, n_3^{te})$ | 96.1 | 43.3 | 59.8 | 70.8 | 64.8 |
| | $(n_2^{tr}, n_2^{tr})$ | $(n_2^{te}, n_3^{te})$ | 95.6 | 37.0 | 53.4 | 67.6 | |

Table D.8: Assessment of the performance of best CNN models in identifying cricket individuals from raw, duration-corrected, and duration- and frequency-corrected five-syllable chirps of new individuals within an open population.

| Data | Train | Test | Precision | Recall | F1 score | Accuracy | Average accuracy |
|---|---|---|---|---|---|---|---|
| Single-night models | | | | | | | |
| Raw | $(n_1^{tr}, n_1^{tr})$ | $(n_1^{te}, n_1^{te})$ | 96.4 | 97.3 | 96.8 | 96.8 | **93.4** |
| | | $(n_2^{te}, n_2^{te})$ | 88.3 | 95.5 | 91.8 | 91.4 | |
| | | $(n_3^{te}, n_3^{te})$ | 88.3 | 96.9 | 92.4 | 92.0 | |
| | | $(n_1^{te}, n_2^{te})$ | 86.7 | 43.4 | 57.8 | 68.4 | **69.1** |
| | | $(n_1^{te}, n_3^{te})$ | 80.9 | 26.9 | 40.3 | 60.3 | |
| | | $(n_2^{te}, n_3^{te})$ | 94.2 | 61.1 | 74.2 | 78.7 | |
| Duration corrected | $(n_1^{tr}, n_1^{tr})$ | $(n_1^{te}, n_1^{te})$ | 93.8 | 96.8 | 95.3 | 95.2 | 89.9 |
| | | $(n_2^{te}, n_2^{te})$ | 86.1 | 97.2 | 91.3 | 90.7 | |
| | | $(n_3^{te}, n_3^{te})$ | 75.9 | 99.4 | 86.0 | 83.9 | |
| | | $(n_1^{te}, n_2^{te})$ | 82.8 | 36.9 | 51.1 | 64.6 | 68.7 |
| | | $(n_1^{te}, n_3^{te})$ | 98.7 | 38.9 | 55.8 | 69.2 | |
| | | $(n_2^{te}, n_3^{te})$ | 92.9 | 48.4 | 63.7 | 72.4 | |
| Duration and frequency corrected | $(n_1^{tr}, n_1^{tr})$ | $(n_1^{te}, n_1^{te})$ | 96.7 | 92.2 | 94.4 | 94.6 | 92.3 |
| | | $(n_2^{te}, n_2^{te})$ | 90.4 | 93.2 | 91.8 | 91.7 | |
| | | $(n_3^{te}, n_3^{te})$ | 89.9 | 97.0 | 91.1 | 90.6 | |
| | | $(n_1^{te}, n_2^{te})$ | 89.5 | 20.6 | 33.5 | 59.1 | 60.9 |
| | | $(n_1^{te}, n_3^{te})$ | 98.7 | 25.1 | 40.0 | 62.4 | |
| | | $(n_2^{te}, n_3^{te})$ | 89.2 | 25.7 | 40.0 | 61.3 | |
| Two-night models | | | | | | | |
| Raw | $(n_1^{tr}, n_1^{tr})$ | $(n_1^{te}, n_1^{te})$ | 98.4 | 98.2 | 98.3 | 98.3 | **95.8** |
| | | $(n_2^{te}, n_2^{te})$ | 96.1 | 98.3 | 97.2 | 97.2 | |
| | $(n_1^{tr}, n_2^{tr})$ | $(n_3^{te}, n_3^{te})$ | 91.4 | 92.8 | 92.0 | 92.0 | |
| | | $(n_1^{te}, n_2^{te})$ | 95.5 | 98.2 | 96.8 | 96.8 | **83.6** |
| | | $(n_1^{te}, n_3^{te})$ | 99.2 | 47.8 | 64.5 | 73.7 | |
| | $(n_2^{tr}, n_2^{tr})$ | $(n_2^{te}, n_3^{te})$ | 99.1 | 61.4 | 75.8 | 80.4 | |
| Duration corrected | $(n_1^{tr}, n_1^{tr})$ | $(n_1^{te}, n_1^{te})$ | 97.5 | 98.4 | 97.9 | 97.9 | **95.8** |
| | | $(n_2^{te}, n_2^{te})$ | 96.6 | 98.7 | 97.6 | 97.6 | |
| | $(n_1^{tr}, n_2^{tr})$ | $(n_3^{te}, n_3^{te})$ | 89.9 | 94.9 | 92.3 | 92.1 | |
| | | $(n_1^{te}, n_2^{te})$ | 97.5 | 97.8 | 97.6 | 96.6 | 83.0 |
| | | $(n_1^{te}, n_3^{te})$ | 99.9 | 52.7 | 69.0 | 76.3 | |
| | $(n_2^{tr}, n_2^{tr})$ | $(n_2^{te}, n_3^{te})$ | 99.9 | 52.5 | 68.9 | 76.2 | |
| Duration and frequency corrected | $(n_1^{tr}, n_1^{tr})$ | $(n_1^{te}, n_1^{te})$ | 97.7 | 97.0 | 97.4 | 97.4 | 95.0 |
| | | $(n_2^{te}, n_2^{te})$ | 97.9 | 96.8 | 97.3 | 97.4 | |
| | $(n_1^{tr}, n_2^{tr})$ | $(n_3^{te}, n_3^{te})$ | 91.3 | 89.1 | 90.1 | 90.3 | |
| | | $(n_1^{te}, n_2^{te})$ | 97.4 | 96.2 | 96.8 | 96.8 | 81.6 |
| | | $(n_1^{te}, n_3^{te})$ | 99.0 | 42.3 | 59.3 | 70.9 | |
| | $(n_2^{tr}, n_2^{tr})$ | $(n_2^{te}, n_3^{te})$ | 99.2 | 54.5 | 70.4 | 77.0 | |

Table D.9: Assessment of the performance of best CNN models in identifying cricket individuals from raw, duration-corrected, and duration- and frequency-corrected five-syllable chirps of known individuals within an open population.
